# Disulfide-bond-induced structural frustration and dynamic disorder in a peroxiredoxin from MAS NMR

**DOI:** 10.1101/2023.02.08.527717

**Authors:** Laura Troussicot, Alicia Vallet, Mikael Molin, Björn M. Burmann, Paul Schanda

## Abstract

Disulfide bond formation is fundamentally important for protein structure, and constitutes a key mechanism by which cells regulate the intracellular oxidation state. Peroxiredoxins (PRDXs) eliminate reactive oxygen species such as hydrogen peroxide by using a catalytic cycle of Cys oxidation and reduction. High molecular-weight assemblies of PRDXs have recently been shown to additionally act as molecular chaperones. The consequences of disulfide bonds on the dynamics of these large assemblies are poorly understood. We show that formation of disulfide bonds along the catalytic cycle induces extensive μs time scale dynamics, as monitored by magic-angle spinning NMR of the 216 kDa-large Tsa1 decameric assembly and solution-NMR of a designed dimeric mutant. We ascribe the conformational dynamics to structural frustration, resulting from conflicts between the disulfide-constrained reduction of mobility and the desire to fulfil other favorable contacts.

## Introduction

Peroxiredoxin enzymes (PRDXs), present in all kingdoms of life, play a key role in detoxifying cells from a variety of peroxides(1–4), and act as peroxide sensors and protein redox regulation factors (5). Tsa1 is a typical 2-Cys peroxiredoxin from yeast, with two conserved cysteines that participate in the catalytic cycle: the peroxidatic cysteine (C_P_, Cys47) is responsible for the catalytic activity by reacting with a peroxide, ROOH, to produce the reduced, non-toxic form, ROH, thereby getting itself oxidised to a sulfenic acid, C_P_–SOH. To regenerate the peroxidatic cysteine, the so-called resolving cysteine (C_R_, Cys170 in Tsa1) forms a disulfide bond with the oxidized C_P_, thereby releasing a water molecule. The disulfide, C_P_–S-S–C_R_, is reduced to the sulfhydryl form, C_P_–SH and C_R_–SH, by the NADPH-dependent Trx/TrxR system (4). During this cycle, the protein undergoes a number of structural modifications, which have been investigated by crystallography. Structures of oxidised and reduced forms, enabled in some cases by mutation of the peroxidatic cysteine, have been reported for several PRDXs (reviewed in Refs. (4, 6)). However, in crystal structures of oxidised states, large parts (>20 residues) have not been modeled, which has hampered the assessment of the structural and dynamical consequences of disulfide bond formation (7, 8). For Tsa1 only the structure of a mutant has been reported, in which the catalytic cysteine C_P_ is mutated to a serine (C47S), and which therefore cannot undergo the functional cycle. PRDXs switch between different oligomeric states (dimers, decamers and higher oligomerisation states) depending on redox state, pH and other factors. The dimer-decamer transition has been observed even directly in cells (9). The dimer-decamer equilibrium is a complex and not entirely understood function of the above-mentioned parameters. Most of the available crystal structures report decameric rings, formed by the assembly of five dimers.

Exciting discoveries over the last years have brought to light an additional role of PRDXs: they can act as chaperones, preventing other proteins from aggregation. Jang *et al*. demonstrated that cytosolic Tsa1/Tsa2 can reversibly assemble to high-molecular weight (HMW) species *in vivo*, and that the HMW species of Tsa1 inhibit aggregation of citrate synthase three times more efficiently than the small heat-shock protein αB-crystallin (10). The conversion to high-molecular weight species of cytosolic Tsa1/Tsa2 in *vivo* is favored by exposing the cell to oxidative stress and heat shock, and thought to be due to oxidation of the catalytic cysteine thiol into a sulfinic acid, Cys–SO(OH) (10–12). Additional factors, including phosphorylation (Thr90 in hPrxI) (13) or exposure to low pH (14), have been shown to induce a functional switch of PRDXs from peroxidases to chaperones. Since these seminal discoveries of a chaperone function, a similar behavior has been found in other 2-Cys PRDXs in different organisms such as SmPrx1 from the parasite *Schistosoma mansoni* (15, 16), the human hPrx1 (13) and the mitochondrial PRDX from *Leishmania infantum* mTXNPx (17–19). The ATP-independent chaperone activity was generally found *in vitro* by biochemical assays; for example, mTXNPx prevents aggregation of citrate synthase at elevated temperatures (18). Understanding the structural and dynamical consequences of disulfide formation and of oligomerisation appears necessary to understand peroxidase function as well as the intriguing chaperone activity of PRDXs.

In this study we use magic-angle spinning (MAS) nuclear magnetic resonance (NMR) spectroscopy to probe dynamics of decameric Tsa1 in its reduced and oxidised forms. Recently developed NEar-rotary Resonance Relaxation Dispersion (NERRD) experiments demonstrate that disulfide formation induces extensive microsecond (μs) motions. Moreover, Bloch-McConnell R_1ρ_ relaxation dispersion MAS NMR reveals μs dynamics in the dimer-dimer interface. A mutation in this interface converts the protein to its dimeric form. Solution-NMR of this dimeric variant shows disorder in the vicinity of the cysteines, suggesting that the dynamic patch observed in oxidised decameric Tsa1 is at least in part present already in the dimer. We find a striking coincidence of the μs dynamics and structural frustration (20), and propose that the induced dynamics in the oxidised form is important for the functional catalytic cycle, and possibly for chaperone function.

## Results and Discussion

### MAS NMR reveals that decameric reduced Tsa1 is overall rigid

We have performed size-exclusion chromatography coupled to multi-angle light scattering (SEC-MALS) and found that at ambient temperature Tsa1 exists primarily in a decameric state, accounting for more than 80% of the total protein concentration (216 kDa; Supplementary Figure S1A). When re-injecting the decameric species into SEC-MALS, only the decameric state is obtained (Fig. S1B), which establishes that we can obtain stable homogeneous decamer samples. We have re-injected the decamer also in the presence of a reducing agent (tris(2-carboxyethyl)phosphine; TCEP) as well as after treating the sample with H_2_O_2_, and found that under all these conditions Tsa1 forms a decameric assembly of ca. 216 kDa (Fig. S1). Based on the slightly altered elution profile upon TCEP-addition leading to two distinct peaks representing the oxidised and reduced decameric species (Fig. S1B), we conclude that the recombinant expressed Tsa1 is in the oxidized state.

NMR is ideally suited to probe dynamics and local conformation at the level of individual atoms, but in solution the NMR signals of particles as large as decameric Tsa1 are broadened due to the associated slow overall tumbling. Indeed, a solution-NMR ^1^H-^15^N spectrum of Tsa1 at ambient temperature shows only very few peaks, even with a deuterated sample and TROSY experiments (23) (Fig. S2). The ^1^H-^15^N peaks observed at ambient temperature correspond to the flexible loop regions, according to the chemical-shift assignment (discussed below).

Magic-angle spinning (MAS) NMR avoids the signal loss/broadening induced by the slow tumbling in solution, by immobilising the proteins (24): in the present case, we obtained MAS NMR samples by sedimenting (ultracentrifuging) decameric Tsa1 from a solution into an MAS rotor (25), thereby retaining a fully hydrated sample (ca. 50% of the rotor volume is buffer). Due to the absence of the overall tumbling in the sediment, MAS NMR spectra are not impacted by the particle size. We used MAS NMR to probe structure and dynamics of the intact decameric assembly at ambient temperature. Under reducing conditions, i. e. in the presence of 5 mM TCEP in the solution, the MAS NMR spectra are of very high quality, as exemplified with a ^1^H-^15^N spectrum in Fig. 1A. The reduced state stayed stable for more than one month in the MAS NMR rotor, as evidenced by unchanged spectra over at least this time period. Using a set of ^1^H-detected 3D H-N-C and H-N-N correlation spectra, we were able to assign the resonances of the majority (72%) of the backbone of this reduced state, Tsa1^red^. The secondary structures, obtained from the chemical-shift assignment and TALOS-N (21), are in very good agreement with the crystal structure of Tsa1^C47S^ that cannot form the (Cys47-Cys170) disulfide bond (Figs. 1B, S3 and S4).

**Fig. 1.**
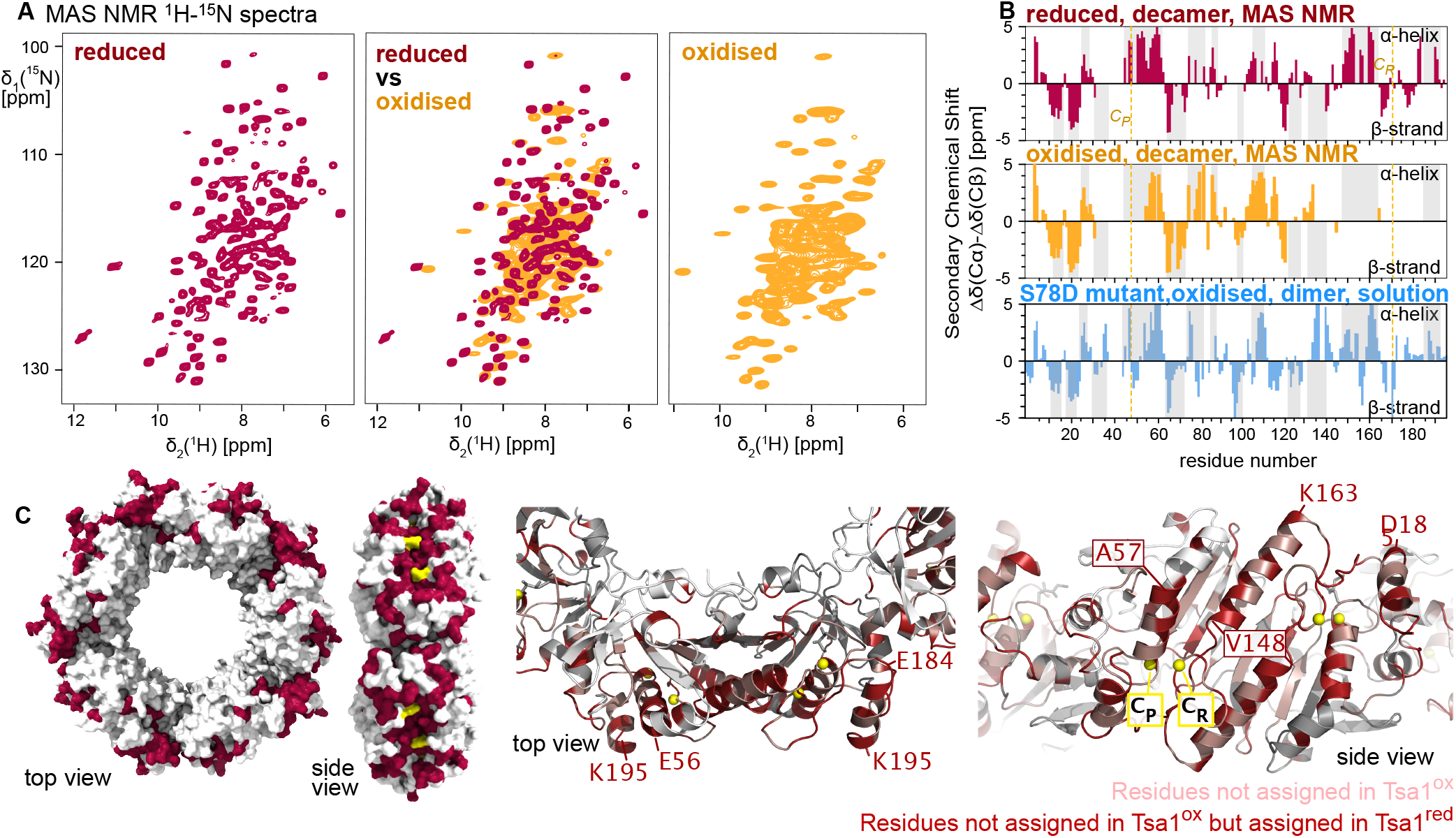
(A)^1^H-^15^N MAS NMR spectra of the native oxidised state, Tsa1^ox^ (orange), and the reduced state, Tsa1^red^ (red) in the presence of reducing agent (TCEP) (^2^H, ^13^C,^15^N labeled protein, MAS frequency of 55.5 kHz, 600 MHz ^1^H Larmor frequency). (B) Secondary chemical shifts that report on residue-wise secondary structure propensities of the decameric Tsa1^ox^ and Tsa1^red^, and the dimeric S78D mutant (from solution NMR, see below). The secondary structures obtained from these data with the program TALOS-N (21) are shown in Fig. S4. Grey bars indicate the secondary-structure elements from the crystal structure of the decameric C47S mutant (3SBC)(22). (C) Location of residues which cannot be detected in Tsa1^ox^. The C_R_ are highlighted in yellow.

### Extensive μs motion in decameric Tsa1 in the oxidised state

The oxidized state, Tsa1^ox^, differs very markedly from the reduced one, as immediately visible from its ^1^H-^15^N spectrum (Fig. 1A). The oxidized state is obtained after purification in non-reducing (native) conditions or, equally, by treatment with H_2_O_2_ (see Methods). Because formation of a disulfide bond links the catalytic cysteine, CP, of one subunit to the CR of another subunit, the presence of a disulfide bond can readily be monitored by the appearance of dimeric species in sodium dodecyl sulfate–polyacrylamide gel electrophoresis (SDS-PAGE). Without any reducing treatment, Tsa1 migrates indeed as a dimer, and upon treatment with dithiothreitol (DTT) only monomeric species are observed (Fig. S5). We have verified by NMR that the native state of a dimeric Tsa1 described below before and after H_2_O_2_ treatment are identical (Fig. S6). The NMR line widths in the decameric Tsa1^ox^ are substantially larger than those of Tsa1^red^ and, importantly, the number of observed cross-peaks is strongly reduced compared to Tsa1^red^: we observed ca. 92 ^1^H-^15^N cross peaks (combining information from 2D hNH and 3D hCANH/hCONH spectra, disregarding side chains peaks), for an expected 184 non-proline residues.

We have considered the possibility that peaks are missing because they are highly flexible and dynamically disordered on ps-ns time scales. If this was the case, the dipolar coupling would be strongly reduced; as the dipolar coupling is the basis for coherence transfer in the above-mentioned experiments, such motion would indeed render transfer inefficient. However the J-coupling is independent of such motion, and J-coupling based experiments may allow transfer, if the motion is fast enough (ps-ns). We observed for Tsa1^ox^ that J-coupling based experiments do not reveal any additional peaks (Fig. S7). This observation allows us to exclude the hypothesis that fast (ps-ns) large-amplitude dynamics, reminiscent of random-coil behavior, causes the loss of those signals that are observed in Tsa1^red^ but not Tsa1^ox^.

We have used dipolar-coupling based 3D and 4D correlation experiments to obtain residue-specific assignments of the observed cross-peaks and were able to assign the vast majority of the detectable backbone resonances, which correspond to 88 residues along the sequence (45%). The near-complete assignment of the peaks observed in the spectrum allows identifying which parts of the structure are not detectable in this oxidised decameric state: the undetected parts cluster around the α-helix that harbors the peroxidatic Cys (residues 38-55), the internal β-strands β6 and β7 (residues 123-130 and 134-140), and the structurally adjacent large C-terminal part (residues 166-196) that comprises a long helix α6 (residues 149-165), a loop region that harbors the resolving-cysteine CR and the C-terminal helix. When seen in the context of the decameric ring, the unobserved residues in Tsa1^ox^ are located on the outer rim of the ring (Fig. 1C).

To identify the origin of the peak loss, we measured ^15^N spin relaxation in Tsa1^ox^ and Tsa1^red^ by MAS NMR. Longitudinal relaxation (R_1_) is sensitive mostly to amplitudes and time scales of motions occurring on ps-ns time scales. Relaxation of ^15^N coherence in the presence of a spin-lock pulse (R_1ρ_) is mostly sensitive to motions on time scales of tens of nanoseconds to hundreds of μs (26–32). Fig. 2A, B shows calculated relaxation rate constants for motion occurring on different time scales. R_1ρ_ experiments can be applied at multiple radiofrequency (RF) spin-lock field strengths, and the dependence on the RF field strength reveals specifically μs-ms motion (Fig. 2B), an effect sometimes referred to as NEar Rotary-resonance Relaxation Dispersion (NERRD; see below) (28, 33, 34). Thus, these different relaxation measurements allow the identification and quantitative interpretation of motional time scales and amplitudes in different time windows.

**Fig. 2.**
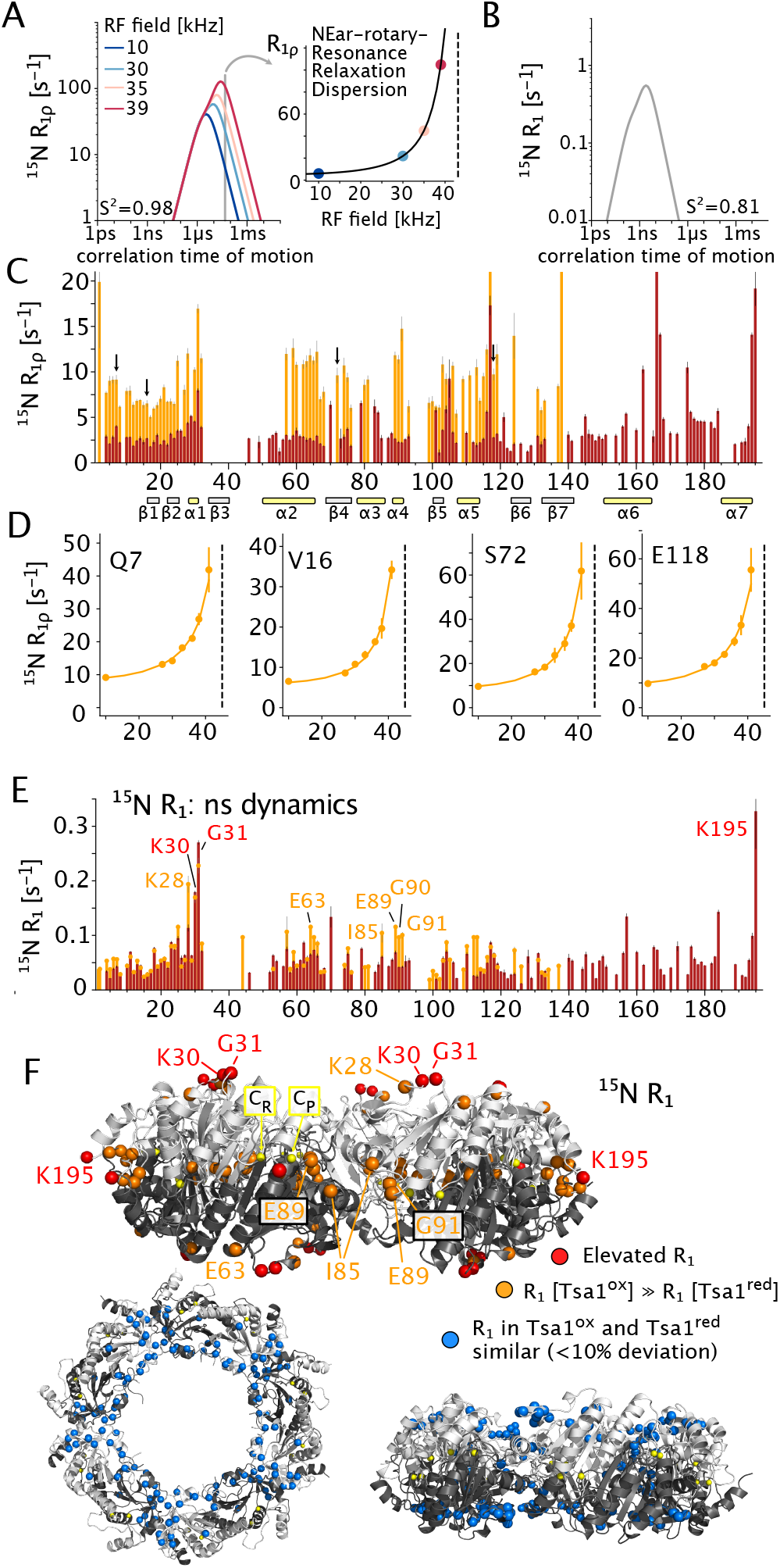
Quantitative dynamics measurements of Tsa1^ox^ (orange) and Tsa1^red^ (red). (A, B) Calculated ^15^N relaxation rate constants for motion of the H-N bond with an amplitude (1-S^2^, where S^2^=1 for a fully rigid site) and a correlation time as shown along the x axis. (A) ^15^N R_1ρ_ relaxation is most sensitive to motion on time scales from hundreds of ns to tens of ms. If the motion is slower than ca. 1 μs, R_1ρ_ depends on the spin-lock field strength, giving rise to NERRD dispersion profiles, i.e. a nonflat profile of R_1ρ_ vs. spin-lock (insert). (B) Calculated longitudinal (R_1_) relaxation rate constants. (C) Experimental R_1ρ_ rate constants in Tsa1^ox^ (orange) and Tsa1^red^ (red), showing strongly enhanced relaxation in the oxidised state across the protein. Average values: R_1ρ_ (Tsa1^red^)= 3.9 ±3.2 s^−1^; 3.3 s^−1^ for residues 1-140 and 5.6 s^−1^ for residues 140–C-terminus. R_1ρ_ (Tsa1^ox^)=9.6 ±4.0 s^−1^. Arrows indicate residues shown in (D). The rectangles at the bottom indicate the positions of β-strands (grey) and α-helices (yellow). (D) Non-flat NERRD profiles unambiguously reveal motion occurring on a μs time scale. (E) Longitudinal ^15^N relaxation in the two states showing that ns motion is similar for the two states for most residues. Residues with enhanced R_1_ relaxation in Tsa1^ox^ compared to Tsa1^red^ are highlighted orange. Residues with particularly high R_1_ are marked in red. Average values: R_1_ (Tsa1^red^)= 0.057 ±0.041 s^−1^. (Tsa1^ox^)=0.063 ±0.037 s^−1^. (F) Visualisation on the 3D structure of the residues with high R_1_ (red), residues with larger R_1_ in Tsa1^ox^ (orange), and residues with similar R_1_ in both states (blue).

In the reduced state, Tsa1^red^, R_1ρ_ rate constants at 10 kHz spin-lock amplitude are ca. 2.9 s^−1^ (median), and longitudinal relaxation rate constants (^15^N R_1_) are ca. 0.05 s^−1^. These are typical values for compact folded proteins without large-scale ns-μs motion (35–41). Residues with higher-than-average relaxation rate constants include K30, G31,E117,V167, which are all located in loop regions, as well as the very C-terminal residues. The larger-amplitude motions reflected by these data is also typical for loops and termini. Overall, the ^15^N relaxation data show that Tsa1^red^ is rather rigid on time scales from ps to tens of μs.

The oxidised state, Tsa1^ox^, differs strongly from its reduced counterpart, in particular with respect to transverse relaxation: the average R_1ρ_ is ca. 3-fold higher, unambiguously showing the presence of motion that is either of larger amplitude or shifted towards the μs range, where R_1ρ_ is highest (Fig. 2A). Longitudinal relaxation, reflecting faster motion, is more similar in the two states. However, the residues for which significantly faster R_1_ relaxation is observed are located in the vicinity of the two cysteines, suggesting that disulfide bond formation enhances not only μs motion but also faster (ns) dynamics (Fig. 2F). Residues further away from the disulfide-bonded parts tend to have more similar R_1_ values (blue in Fig. 2F). The overall enhanced transverse relaxation in Tsa1^ox^ also provides a rationale why J-coupling based experiments do not provide additional (but rather much less) signal, as discussed above.

To obtain more precise information about this motion, we performed ^15^N R_1ρ_ NEar-rotary Resonance Relaxation Dispersion (NERRD) experiments, i.e. we measured R_1ρ_ rate constants as a function of the applied spin-lock field strength, approaching the regime where the spin-lock field strength (nutation frequency ν_RF_) approaches the MAS frequency ν_MAS_. Non-flat NERRD profiles directly point to motion on time scales of μs (see Fig. 2A). The underlying mechanism is the fluctuation of the H-N bond orientation (26). In Tsa1^ox^ we observe strong NERRD effects for all observed residues (Fig. 2D and S8). To gain more quantitative insight, we jointly fitted eight relaxation rate constants (seven R_1ρ_ and one R_1_) with the DETECTORS approach (42). This formalism reports on the amplitudes of motions in different time windows (named detectors). Rather than fitting μs dyamics as a single process at one time scale, as commonly done in solution and solids (43), the spin relaxation is modeled as resulting from combined dynamical processes with correlation times across the ns-ms range (Fig. S9). This analysis points, on the one hand, to enhanced ns motion of the loop region comprising K30 and G31 (revealed by the R_1_ data of Fig. 2E, F). On the other hand, μs motion is found for most residues across the protein in Tsa1^ox^, mirroring the generally high levels of R_1ρ_ and the non-flat NERRD profiles. The last observed residue, T138, as well as residues around the nonassigned part (residues 34–56), have particularly enhanced μs mobility; this is also evident from the data at 10 kHz RF field strength (Fig. 2C). This finding supports the notion that the parts that are not visible in Tsa1^ox^ are broadened due to even larger-amplitude μs motion.

We have furthermore applied Bloch-McConnell relaxation dispersion (BMRD) experiments, which detect μs-ms exchange processes based on chemical-shift fluctuations, rather than bond-angle fluctuations (which are the basis of NERRD): in BMRD, μs–ms modulation of the chemical shift leads to enhanced transverse relaxation, which can be quenched by a spin-lock that is sufficiently strong; thus, a decrease of R_1ρ_ rate constants when increasing the RF field strength from ca. 2 to above 10 kHz reveals chemical-shift fluctuations on μs-ms time scales (28, 33, 43, 44). Below ca. 2 kHz, insufficiently suppressed dipolar dephasing may lead to increased R_1ρ_ (33).

Figure 3 shows that residues with significant BMRD effects are located at the interface of dimeric building blocks. Interestingly, similar effects are found for both the oxidised and reduced species. We ascribe this process to some flexibility within the dimer-to-dimer interface, which prompted us to investigate more closely the dimeric state and the oligomeri-sation.

**Fig. 3.**
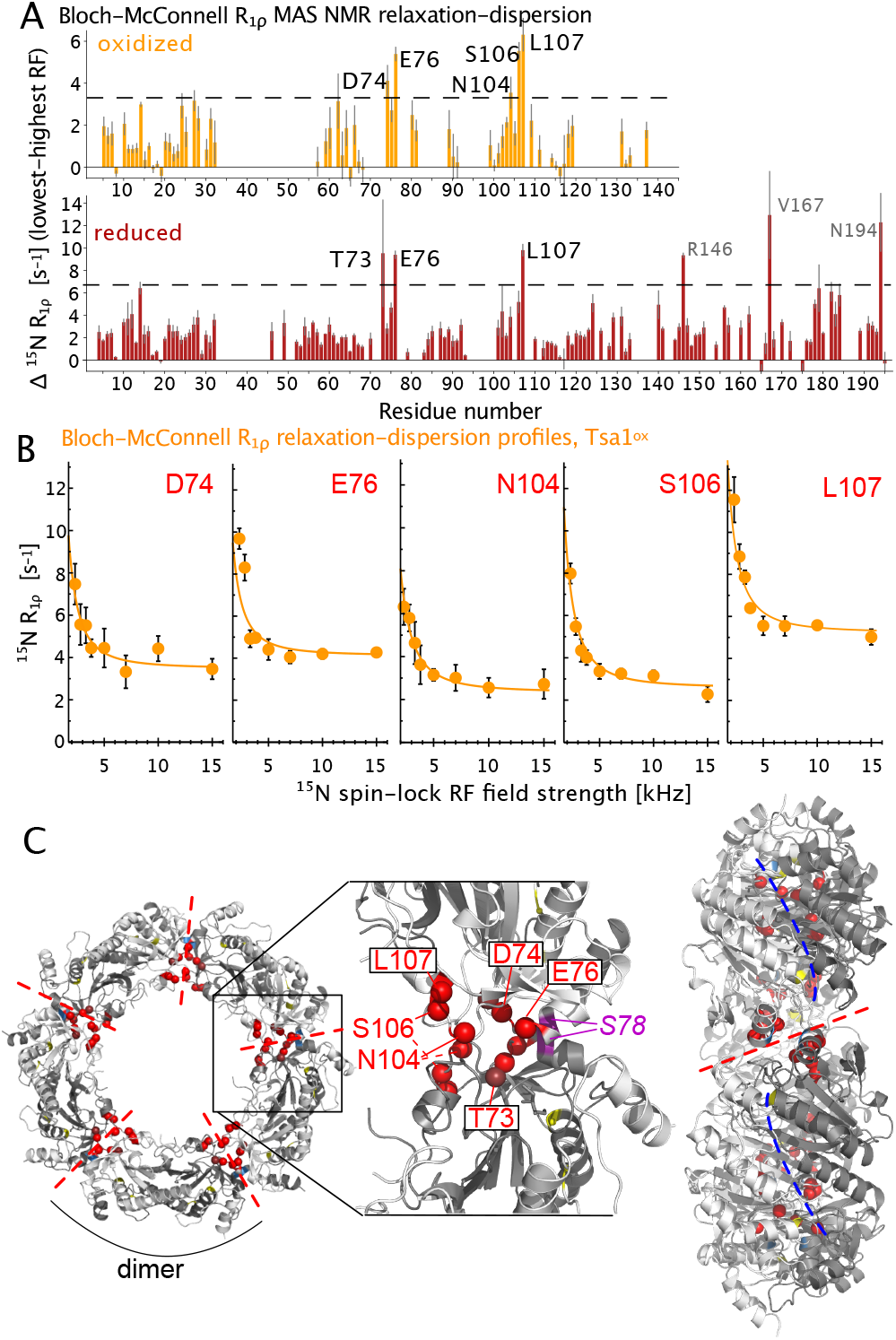
Bloch-McConnell R_1ρ_ MAS NMR relaxation-dispersion (BMRD) data of Tsa1^ox^ (orange) and Tsa1^red^ (red). (A) Difference of R_1ρ_ obtained at low RF field strength (2.3 kHz) minus R_1ρ_ at high RF field strength (15 kHz for Tsa1^ox^; 10 kHz for Tsa1^red^). The horizontal line indicates the three-fold standard deviation over all residues. (B) BMRD profiles for Tsa1^ox^ for the residues with the largest dispersions. (BMRD curves of all residues are shown in Fig. S11.) The R_1ρ_ rate constants have been corrected for the NERRD effect, by back-calculating and subtracting the rate constants based on DETECTORS fits (42) of the NERRD data of Fig. 2D, as outlined in Fig. S10). Residue-wise frequency offsets have been corrected with R_1_ data, as commonly done (43). Solid-lines show a joint fit of the 5 residues (k_ex_=1100 s^−1^) to a two-state exchange model. (C) Location of the residues with the largest BMRD effects in the decameric Tsa1 structure (PDB 3SBC).

### Solution-NMR of dimeric Tsa1 reveals disorder around the cysteines

To understand the effects of decamer-formation from dimers on structural and dynamical properties, we searched for ways of stabilising the dimer. At ambient conditions Tsa1 predominantly exists in the decameric state, where solution-NMR spectra are of poor quality (Figs. S1, S2, S12). We followed how temperature changes the oligomerisation state by performing methyl-detected diffusion-ordered NMR spectroscopy (DOSY) in solution using a deuterated, Ile/Leu/Val methyl-labeled sample. Because of the high intrinsic sensitivity of methyl groups, together with the methyl-TROSY effect (45), the signals of methyl groups are detectable even in the 216 kDa-large decameric Tsa1 assembly (Fig. S12), which allowed us to quantify the diffusion coefficient. Temperature-dependent DOSY data reveal the transition from the decameric to a dimeric state with a mid-point at ca. 315 K above which Tsa1 is predominantly dimeric.

However, prolonged experiments at such high temperatures are not compatible with the sample integrity, which prompted us to insert a mutation in the dimer-dimer interface that disrupts decamer formation also at low temperature. The S78D mutant, identified before (46), appears entirely dimeric as seen by size-exclusion chromatography (Fig. S1C). Note that S78 is located close to the cluster of residues for which we detected μs motion in the decameric state by MAS NMR (Fig. 3C). In contrast to the wild-type protein, Tsa1^S78D^ yields excellent solution-NMR spectra of both backbone and methyl groups already at 298 K (Fig. S2 and S12). We assigned the backbone resonances as well as a large extent of the methyl resonances of Ile (δ1), Val (γ1 and γ2) and Leu (δ1 and δ2) of the oxidised state, which corresponds to the form obtained after purification from *E. coli* (Fig. S6).

The chemical-shift derived secondary structure propensity of the S78D dimer is largely in agreement with the crystal structure of Tsa1 and with the chemical-shift derived secondary structures of the decameric state by MAS NMR, but there are notable exceptions (Fig. 1B). Importantly, the part that carries the peroxidatic Cys-47 (α2) does not have chemical shifts of a stable α-helix, unlike in the decameric state, which shows that this helix is marginally stable in the Tsa1^S78D^ dimer.

To gain direct insights into the dynamics, we have performed backbone ^15^N amide and Ile, Leu, Val methyl ^1^H-^13^C triple-quantum relaxation experiments (Fig. S12 and S14). Backbone ^15^N transverse relaxation rate constants (R_1ρ_) are ca. 2-3 fold reduced for residues located C-terminal to residue 180, including the part the forms helix α7 in the decamer. Moreover, the methyl order parameters of residues in this part (V167, L168, I179, V183) have low order parameters (S^2^~0.1), much lower than for the majority of the protein, providing independent evidence that the C-terminal part has significant disorder in the dimeric, oxidised Tsa1^S78D^ (Fig. 4). These findings show that the disorder in this part of the protein exists also in the dimer.

**Fig. 4.**
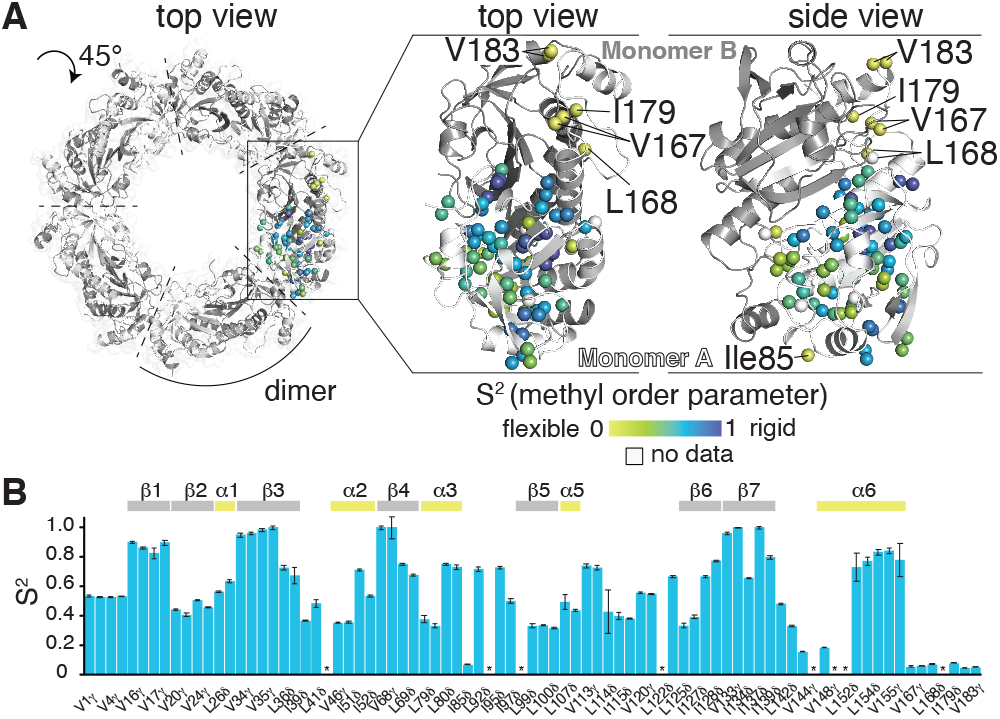
Dynamics of the dimeric state, probed in Tsa1^S78D^ by Ile, Leu and Val methyl order parameters measured at 37°C. (A) Methyl order parameters (S^2^) plotted on the structure of one Tsa1 monomer as indicated by the yellow to blue gradient. The top view was rotated by 45° along the central decamer axis compared to Fig. 1C. Residues in the C-terminal part and the surface exposed Ile85, showing the highest degree of dynamics, are annotated. (B) S^2^ values plotted against the sequence. Tsa1 secondary structure elements are indicated on the top. Residues for which no data could be obtained are marked with a *.

### Structural frustration around the dynamic disulfide part

Collectively, our NMR data have revealed that the formation of the disulfide bond induces dynamic disorder. As a consequence, signals in MAS NMR spectra of the oxidised state are broadened beyond detection, and we find broadening and disorder (in particular of the peroxidatic Cys–(C_P_)-carrying helix and the C-terminal ca. 20 residues) also in the dimer. One may expect that the μs motion that we have revealed here is reflected in structural heterogeneity when viewed by X-ray crystallography. This hypothesis can be tested for the case of the human homolog (PRDX2), for which structures of reduced (C_P_–SH), hyperoxidised (C_P_–SO_2_H) (47) and oxidised (C_P_–S-S–C_R_) (7) decameric states have been determined. For the reduced and hyperoxidised states of PRDX2, as well as for reduced-state-mimicking Tsa1^C47S^, essentially the entire protein has been modeled at ca. 1.7 to 2.3 Å resolution, and with only modestly increased B-factors towards the very C-terminal residues and low B-factors for the C_P_-carrying helix (Supplementary Fig. S16). In the oxidised state, however, the entire C-terminal part, comprising the two last helices and the connecting loop, as well as the helix containing the CP have elevated B-factors, and ca. 40 residues have not been modeled in the crystallographic model; moreover, superposition of the ten subunits of the decamer reveals significant structural heterogeneity (Fig. S16E).

We propose that this dynamic disorder induced by the C_P_–S-S–C_R_ bond formation is due to conflicting structural constraints within the protein or structural frustration (20), which arises when multiple energetically favorable interactions cannot be simultaneously fulfilled. As a consequence of structural frustration, the protein explores a broad shallow conformational landscape on longer time scales, rather than populating a narrow range of conformations with low-amplitude (fast) fluctuations. To explore this idea we have used the available crystal structures and determined the structural frustration for the reduced and oxidised states of PRDX2 (Fig. 5A) and also of the reduced-mimicking state of Tsa1 (C47S mutant; no crystal structure of oxidised Tsa1 is available; Fig. 5B). This data shows a cluster of structurally frustrated sites at the outer rim of the decameric ring (ca. residues 165 to the C-terminus) already in the reduced state (red in Fig. 5A and right side in panel B).

**Fig. 5.**
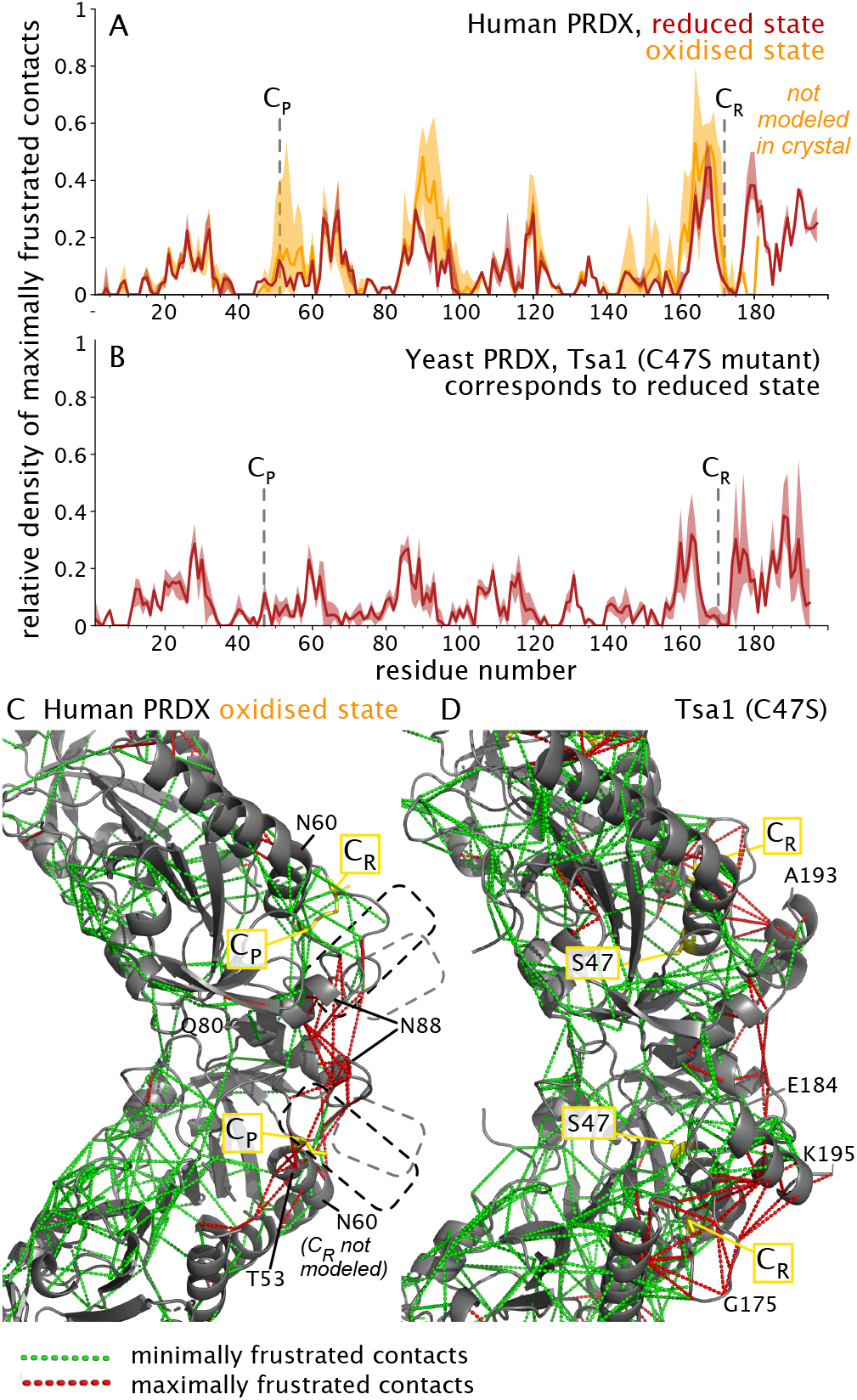
Structurally frustrated parts of PRDX coincide with the outer rim where extensive line broadening is observed. (A, B) Density of maximally frustrated contacts in human PRDX (A) in the oxidised and reduced states (PDB 5IJT (7) and 7KIZ (47), respectively) and the Tsa1 mutant C47S that mimicks the reduced state (PDB 3SBC (22)), plotted along the sequence position. The data have been obtained with the Frustratometer webserver (48, 49). The solid line shows the average value over all available data in the decamer, and the shaded area denotes the range between the highest and lowest value found. (C, D) Clusters of maximally and minimally frustrated contacts, shown by red and green dashed lines, respectively, on the structures of human PRDX^ox^ (C) and Tsa1^C47S^. The data have been obtained in the same Frustratometer analysis as in (A, B). In the structure of PRDX^ox^, some parts have not been modeled (located in regions indicated by dashed boxes; see also Fig. S16), including some of the C_R_ cysteines. Additional data for another homolog are shown in Fig. S15.

The oxidised state has a substantially increased number of maximally frustrated sites which cluster in particular in the parts comprising the two cysteines and the structurally neigh-boring part of residues ca. 80-100. As noted, a large C-terminal stretch that forms an α-helix in the reduced state is not observed in the oxidised state. Taken together, in light of the strong correlation of dynamic (μs motions, blurred electron density) and structurally frustrated sites, we propose that structural frustration induced by disulfide-bond formation is at the core of the functional cycle of PRDXs.

## Conclusion

Disulfide bond formation is generally associated with an increased stability of a folded state (50). Contrary to this view, our quantitative dynamics investigations have shown the appearance of collective μs motions upon formation of the key catalytic disulfide bond in a peroxiredoxin. Our data mirror the lack of visible density in crystallographic structures of the decameric assembly. Our solution-NMR data, using a mutant that stabilises the dimer, point to dynamic disorder already in the dimeric state. Our data reveal a link between slow (μs) dynamics, structural frustration and disulfide-bond formation. Although the extensive dynamics hamper the actual visualisation of the structural ensemble at the atomic level, we propose that the disulfide-induced dynamics likely leads to exposure of hydrophobic patches and increase of the conformational entropy. Both of these effects have been ascribed to chaperone function: the increased entropy would lower the free-energy penalty associated with binding a disordered protein to the chaperone, and the exposure of hydrophobic patches is, generally, a key to chaperone mechanisms (51, 52). Ongoing studies assess the link of the observed structural frustration and chaperone activity of PRDXs.

## Methods

### Cloning

The plasmid harboring wild-type Tsa1 (pET19b-Tsa1) was originally obtained with an amino-terminal deca-histidine-tag in a codon-optimized manner from GenScript (kind gift from P.O. Widlund/T. Nyström). A S78D single point mutation was introduced into the Tsa1 plasmid by standard methods using the following forward 5’ GGT CCA CGC CAG CAG GTC GTA TTC GCT ATC GGT G 3’ and reverse 5’ CAC CGA TAG CGA ATA CGA CCT GCT GGC GTG GAC C 3’ primers. Primers were purchased from Eurofins.

### Protein expression and purification

Tsa1 wild-type or Tsa1^S78D^ plasmids were chemically transformed into E. *coli* BL21(DE3) competent cells. The cell culture was initiated by inoculation of a single colony into a 20 mL pre-culture of LB medium supplemented with 100 μg/ml Ampicillin, and subsequently incubated at 37°C overnight. This pre-culture was used to inoculate 1L of LB culture medium supplemented with 100 μg/ml Ampicillin, and cells were grown until an OD_600_ ~ 0.6 was reached. Expression was induced by the addition of 0.3 mM IPTG and cells were grown for an additional 3 hours at 37°C. Cells were harvested by centrifugation at 6000 g for 25 min at 4°C, subsequently resuspended in lysis buffer (20 mM Tris pH 8, 250 mM NaCl, 5 mM imidazole), and flash-frozen in liquid nitrogen until purification. After thawing the cells, one cOmplete EDTA-free Protease Inhibitor Cocktail tablet (Roche), HL-SAN DNAse I (ArcticZymes) and 1 mM MgCl_2_ were added to the lysate. Cells were lysed by 4 passes through an Avestin Emulsiflex C3 (pressure between 20.000–25.000 psi) and centrifuged at 24.000 g for 45 min at 4°C. The cleared supernatant was applied to a Ni^2+^–HisTrap column (GE Healthcare) and eluted with a 3 steps imidazole gradient in lysis buffer ranging from 150 mM to 500 mM and finally 1 M imidazole. Tsa1 typically eluted within the 500 mM imidazole fraction. The fractions containing Tsa1 were dialyzed against lysis buffer overnight at 4°C, concentrated by ultracentrifugation (Vivaspin concentrators, Sartorius), and applied to a gel filtration column (Superdex S200 increase, GE Healthcare) equilibrated with NMR buffer (typically 50 mM potassium-phosphate, pH 7.4, 50 mM KCl or PBS pH 7.4). Fractions of the same oligomeric state were concentrated and the concentration of the sample was quantified by measuring the optical density (OD_280_) based on the theoretical molar extinction coefficient of 24500 M^−1^ cm^−1^.

### Isotope labelling

Isotope labelled proteins were expressed either in H_2_O or D_2_O based M9 minimal media supplemented with the desired isotopes (53). For the uniformly labeled [U-^15^N] and [U-^15^N, ^13^C] proteins, (^15^NH_4_)Cl and D-(^13^C)-glucose were used, whereas for [U-^2^H,^15^N,^13^C] labelling, D_2_O-based M9 medium supplemented with (^15^NH_4_)Cl and D-(^2^H,^13^C)-glucose were used. For production of specific methyl-group labelled Tsa1, D_2_O-based M9 medium with D-(U-^2^H, ^12^C)-glucose and (^15^NH_4_)Cl was used, and deuterated biosynthetic precursors (keto-acids, see below) with specific ^13^CH_3_ labelling were added to the culture one hour prior to induction at OD_280_ around 0.5: for labeling of isoleucine, leucine and valine (ILV) methyl groups, 85 mg/L of α-ketoisovalerate-(3-methyl-^13^C),4-^13^C,3-d and 50 mg/L of [^2^H], 3,3-[^13^CH_3_]-ketobutyrate were added to the medium resulting in [U-^2^H,^15^N, Ile-δ1-^13^CH_3_, Leu/ Val-^13^CH_3_] labeled Tsa1 and Tsa1^S78D^ (54). The stereospecific labeling of valine methyl groups only was achieved by adding 85 mg/L of α-ketoisovalerate-(3-methyl-^13^C),4-^13^C,3-d and 40 mg/L of L-leucine-d10, to produce [U-^2^H,^15^N, Val-^13^CH_3_] labeled Tsa1^S78D^ sample (55). All stable isotopes were purchased from Merck.

### Solution NMR spectroscopy

NMR measurements were performed on Bruker Avance III HD 700 MHz or 800 MHz spectrometers, using Topspin 3.5 software and equipped with either TCI or TXO cryogenically cooled triple resonance probes. All experiments were performed either in 50 mM KPi, 50 mM KCl, pH 7.4 buffer or PBS pH 7.4 at the indicated temperatures. For the sequence specific backbone resonance assignments of [U-^2^H,^15^N,^13^C] Tsa1^S78D^ the following TROSY-type experiments were recorded at 37°C: 2D [^15^N, ^1^H]-TROSY (23), 3D trHNCO, 3D trHNCA, 3D trHNCACB, 3D trHNCOCACB (56). Complementary sequence specific backbone and side-chain assignments of [U-^15^N, ^13^C] Tsa1^S78D^ were recorded: BEST-type triple resonance experiments 3D BT-HNCA+, 3D BT-HNCO (57), BT-HNCACB+ (57, 58) as well as a CBCA(CO)NH (59). In deuterated proteins, the re-protonation of all exchangeable amide sites is important when using such amide-detected experiments. To verify that no amide hydrogen sites are lost due to the deuterated sample production (in D_2_O), we compared ^1^H-^15^N spectra of protonated and deuterated Tsa1^S78D^ (Supplementary Fig. S17); no additional signals were found in the sample produced in H_2_O. Aliphatic side-chain resonance assignment for [U-^15^N, ^13^C] Tsa1^S78D^ was performed based on 2D ^1^H-^13^C HMQC spectra with/without constant time version, as well as 3D HBHA(CO)NH, and HCCH-TOCSY-experiments (59). To confirm methyl group assignments on ILV-samples 3D ^13^C methyl-SOFAST-NOESY experiments (60, 61) with mixing times of 50 ms and 600 ms as well as a 3D HMBC-HMQC spectrum (62) were measured in 99.8% D_2_O-based NMR buffer. For quantitative analysis of signal intensities, the amplitudes were corrected by differences in the ^1^H-90° pulse-length, the number of scans, and the dilution factor (63). Further, a weighting function with weights 1–2–1 for residues (i-1)–i–(i+1) was applied to the raw data (64). NMR data were processed with a combination of NMRPipe (65) and mddNMR2.6 (66), and analyzed using CARA (67). Secondary chemical shifts were calculated relative to the random coil values calculated by the POTENCI program (68). Methyl group assignment was performed using the MAGIC algorithm (69) and confirmed by a combination of manual analysis of the side-chain TOCSY, HMBC-HMQC as well as NOESY data sets. This approach yielded the following degree of assignment (~ 92%): Ile δ1 (13/13), Leu δ1, δ2 (30/34), and Val γ1, γ2 (34/36).

Translational diffusion coefficients were measured by recording a series of 1D ^13^C-edited DOSY spectra at different temperatures ranging from 25°C to 60°C, using a pulse scheme (^13^C-edited BPP-LED (70)) that is similar to an ^15^N-edited BPP-LED experiment with ^15^N and ^13^C pulses interchanged (71). The gradient duration *δ* was adapted to 3.2 ms instead of 4.8 ms used in the ^15^N-filtered version and a diffusion delay T of 400 ms and a *τ* of 0.1 ms were used. The strength of the encoding/decoding was increased stepwise. The resulting ^1^H signal was integrated over the methyl ^1^H frequency range to obtain intensities as a function of encoding/decoding gradient strength. In addition, 1D ^1^H diffusion experiment (DOSY) with ^13^C filter and decoupling were recorded at the same temperatures. To adjust for the temperature-dependent changes in viscosity of the buffer, the obtained translational diffusion coefficient was adjusted on the basis of temperature-dependent viscosity values for D_2_O-based buffers over the used temperature range (72). DOSY data were compared to previously reported data to obtain a quantitative benchmark (73–75), as described in Fig. S13. Structure-based diffusion coefficients (Fig. S13) were obtained with HYDROPRO (76).

### Magic-angle spinning NMR sample preparation

2.5 mg of [U-^2^H,^15^N,^13^C] Tsa1^WT^ protein sample in 50 mM KPi, 50 mM KCl, pH 7.4 buffer was thawed at room temperature and sedimented into a 1.3 mm rotor using ultracentrifugation at 50000 g overnight at 6°C, using an in-house-built device for ultracentrifuges.

### Magic-angle spinning NMR spectroscopy

MAS NMR experiments were recorded on a Bruker Avance-III spectrometer at a ^1^H Larmor frequency of 600 MHz, equipped with a 1.3 mm probe tuned to ^1^H, ^13^C, ^15^N and ^2^H frequencies. Sequence-specific backbone resonance assignments of [U-^2^H,^15^N,^13^C] Tsa1^WT^, have been recorded at 25°C effective sample temperature at 55 kHz magic angle spinning (MAS) by the following ^1^H detection experiments: 2D hNH, 3D hCONH, 3D hCOcaNH, 3D hCANH, 3D hCAcoNH. 3D hCACBcoNH, 3D hCACBcacoNH, 3D hcaCBCAcoNH, 4D hCACONH, 4D hCOCANH and 4D hCACBcaNH and 3D hNcocaNH and 3D hNcacoNH experiments. The experiments use cross-polarization (CP) transfer steps for all heteronuclear transfers; for CA-CO transfer a BSH-CP was used (77); CA-CB out-and-back transfers used 6.5 ms-long INEPT transfers. All experiments were used as implemented in the NMRlib library (78). The NMR data were processed with NMRPipe (65) and subsequently analyzed with CcpNmr Analysis 3.0.1 (79).

### NMR backbone and side chain dynamics

For the analysis of the dynamic properties of Tsa1^S78D^, the following relaxation experiments were measured: R_1_(^15^N) (80), ^15^N(^1^H)-NOE (hetNOE) (80), R_1ρ_(^15^N) (81), and TROSY for rotational correlation times (TRACT) (82). Non-linear least square fits of relaxation data were done with NMRFAM-Sparky 1.47 (83). Side-chain dynamics experiments were performed on a [U-^2^H,^15^N, Ile-δ1-^13^CH_3_, Leu-, Val-^13^CH_3_] labeled Tsa1^S78D^ sample at 37 °C in 99.9% D_2_O based NMR buffer. Side chain methyl order parameters (S^2^) were determined by a relaxation-violated coherence-transfer triple-quantum experiment (84). The buildup of triple-quantum (3Q) coherence was monitored in a series of ^1^H–^13^C experiments at different delay times (13 values from 1 to 35 ms). The decay of single-quantum (SQ) coherence was followed in a set of reference experiments at the same delays. Ratios of the peak intensities of these two sets of experiments were fitted to obtain of site-specific order parameter S^2^ using the seperately determined (TRACT-approach) overall correlation time τ_c_ in 100% D_2_O of 27 ns.

For the analysis of the dynamic properties of Tsa1^WT^, ^15^N-R_1_ and ^15^N-R_1*ρ*_ relaxation experiments were measured by MAS NMR on the [U-^2^H,^15^N,^13^C] Tsa1^WT^ sample. The experiments were based on hNH 2D experiments with CP transfer (38). ^15^N-R_1_ was measured at a MAS frequency of 55 kHz, and relaxation delays of 0.1 to 15 s. A series of ^15^N-R_1*ρ*_ experiments was collected at delay times ranging from 1 ms to 200 ms at ^15^N spin-lock radio-frequency field strengths from 2.3 to 15 kHz, resulting in a Bloch-McConnell-type relaxation-dispersion experiment (33). This approach monitors microsecond-millisecond dynamics by the chemicalshift fluctuations. NEar-Rotary-Resonance relaxation Dispersion (NERRD) ^15^N-R_1*ρ*_ experiments were recorded at a MAS frequency of 45 kHz, which allows approaching the rotary-resonance condition (spin-lock RF field equals MAS frequency) without requiring extensively high RF fields. A series of 2D spectra with spin-lock durations ranging from 1 ms to 50 ms was recorded at RF field strengths from 10 to 41 kHz. This experiment detects μs-ms motion involving bond reorientation (33, 34). All experiments are implemented in NMRlib (78).

The analysis of the NERRD data was performed using the DETECTORS program, a python-based tool (github.com/alsinmr/pyDR). We used either three or four DETECTORS, both of which give similar qualitative result (see Fig. S9).

Given the large NERRD effects for almost all residues, which extends to even below 10 kHz, the analysis of Bloch-McConnell relaxation-dispersion (BMRD) data involved a correction for the NERRD effect, as outlined in Fig. S10. In brief, the DETECTORS parameters were used to back-calculate R_1ρ_ rate constants at all RF fields used in the BMRD experiment, and subtracted from the experimental rate constants. These rate constants are close to zero. Analysis of the non-flat profiles (shown in Fig. 3) was done with the program relax (85). As the program cannot handle negative relaxation rate constants, which result from the above-described subtraction procedure, a constant value (5 s^−1^) was added to all data. This plateau value is of no influence for the data analysis.

### Preparation of reduced Tsa1 samples

In order to obtain a reduced sample, 2.1 mg of [U-^2^H,^15^N,^13^C] Tsa1^WT^ was incubated at 30 °C for 30 min with 5 mM DTT. The protein solution was quickly centrifuged at 10000 g in a table centrifuge to pellet down eventual aggregates. Protein supernatant was sedimented into a 1.3 mm rotor overnight by ultracentrifugation at 50000 g, 6°C. The protein sample showed a stability in a reduced state for several weeks, as evidenced by 2D hNH MAS NMR experiments.

### Size-exclusion chromatography - multi-angle light scattering

SEC-MALS experiments were performed using a Superdex Increase 200 10/300 GL column (GE Healthcare) on an Agilent 1260 HPLC Infinity II in phosphate buffer (PBS or 50 mM KPi KCl pH 7.4) at RT (ca. 297 K). Protein elution was monitored by three detectors in series namely, an Agilent multi-wavelength absorbance detector (absorbance at 280 nm and 254 nm), a Wyatt miniDAWN TREOS multiangle light scattering (MALS) detector, and a Wyatt Optilab rEX differential refractive index (dRI) detector. The column was pre-equilibrated overnight in running buffer to obtain stable baseline signals from the detectors before data collection. Molar mass, elution concentration, and mass distributions of the samples were calculated using the ASTRA 7.1.3 software (Wyatt Technology). A BSA solution (2–4 mg/ml), purchased from Sigma-Aldrich and directly used without further purification, was used to calibrate inter-detector delay volumes, band broadening corrections, and light-scattering detector normalization using standard protocols within ASTRA 7.1.3.

## Data availability

The NMR chemical shift assignments of Tsa1 in the decameric oxidised state, decameric reduced state by MAS NMR and of Tsa1^S78D^ by solution-state NMR have been deposited in the BioMagResBank (accession number 51788). Other data are available from the authors upon request.

## Acknowledgements

We thank Albert A. Smith (Dresden) for discussions and help with DETECTORS. Intramural funding from Institute of Science and Technology Austria is acknowledged. This work used the platforms of the Grenoble Instruct-ERIC center (ISBG; UMS 3518 CNRS-CEA-UJF-EMBL) within the Grenoble Partnership for Structural Biology (PSB), as well as the Swedish NMR Centre of the University of Gothenburg. Both platforms provided excellent research infrastructures. B. M. B. gratefully acknowledges funding from the Swedish Research Council (Starting grant 2016-04721), the Swedish Cancer Foundation (2019-0415), and the Knut och Alice Wallenberg Foundation through a Wallenberg Academy Fellowship (2016.0163) as well as through the Wallenberg Centre for Molecular and Translational Medicine, University of Gothenburg, Sweden.

## Author contributions

L. T. produced all Tsa1 protein samples, performed and analysed biochemistry experiments, solution- and MAS-NMR experiments. A. V. performed MAS NMR experiments. B. M. B. performed and analysed solution-NMR experiments. P. S. performed and analysed MAS NMR experiments. L. T., B. M. B. and P. S. designed the study. P. S. and L. T. prepared figures and wrote the manuscript, with input from all authors.

---

**Fig. S1.**
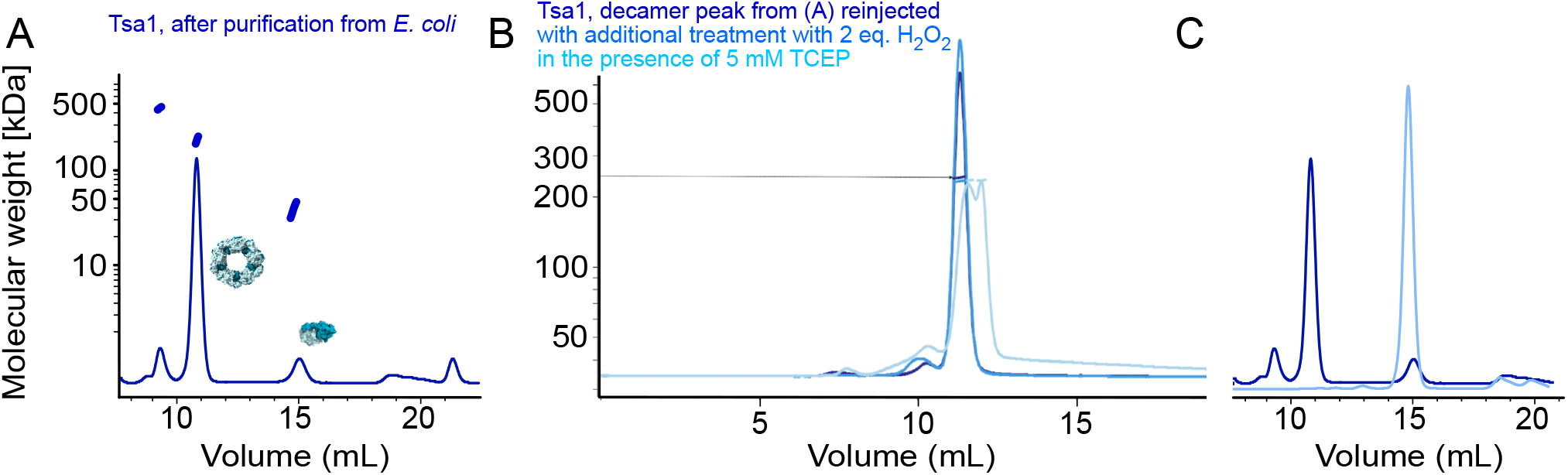
Size-exclusion chromatography coupled to multi-angle light scattering (SEC-MALS) characterisation of wild-type Tsa1, showing the retention volume (x axis) and the absorbance (continuous lines) and the molecular weight obtained from scattering (short lines at the elution peaks). (A) SEC-MALS profile for Tsa1 obtained at the end of the purification of the protein from *E. coli* (see Methods). This data shows that the protein forms predominantly decameric species (ca. 216 kDa), and a small amount of 20-mer and dimer species. (B) Re-injection of the decamer peak from (A) without treatment (dark blue) and with oxidative or reductive treatment. This data shows that the oligomerisation state of the protein is the same in both oxidized and reduced states. (C) SEC profile of the S78D mutant (light blue). For comparison, the profile of the wild-type protein is shown (dark blue).

**Fig. S2.**
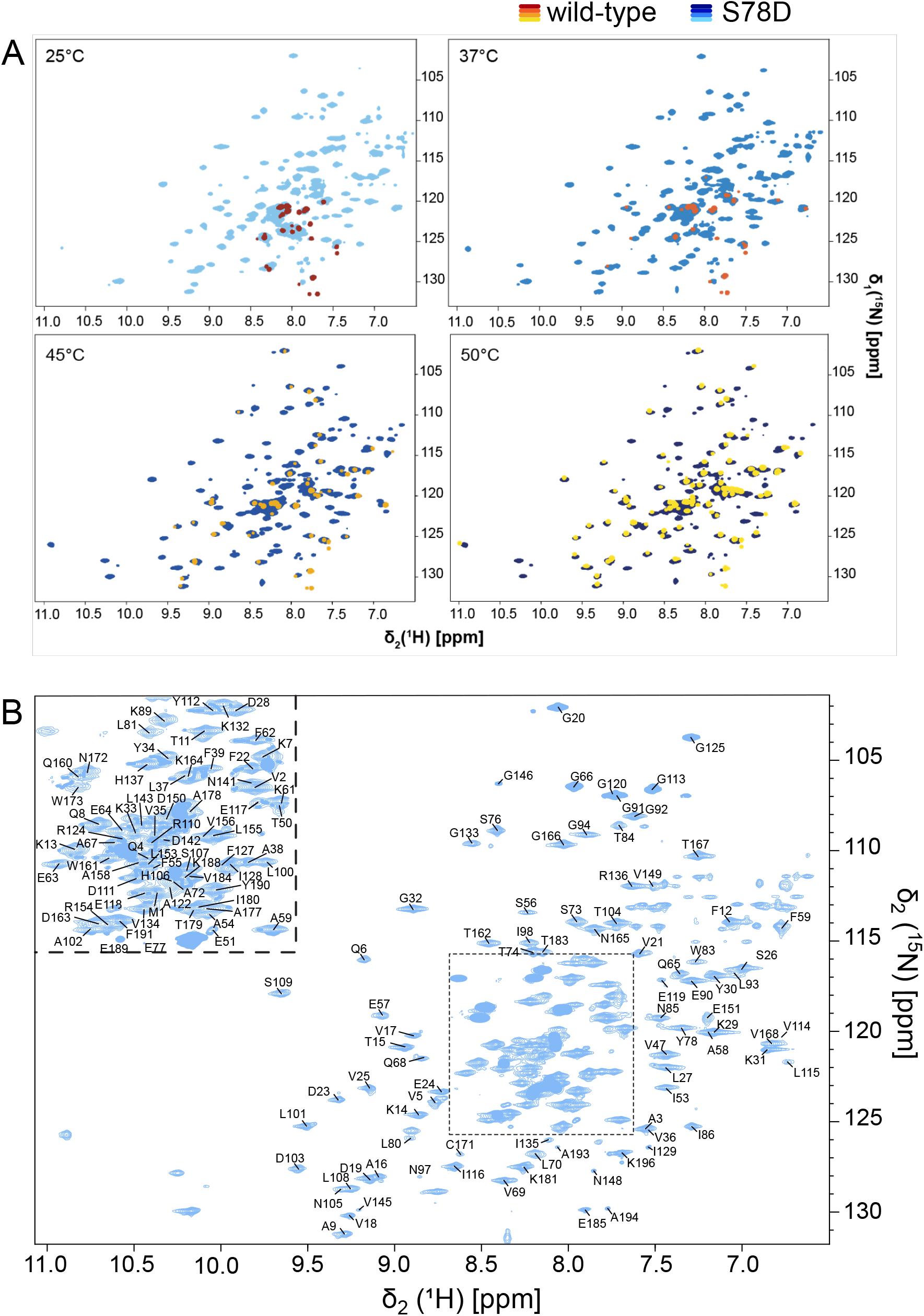
(A) Solution-state ^1^H-^15^N correlation spectra of Tsa1^WT^ and Tsa1^D78D^ at different temperatures, as indicated. (B) Spectrum of Tsa1^D78D^ at 37 °C with sequencespecific resonance assignments.

**Fig. S3.**
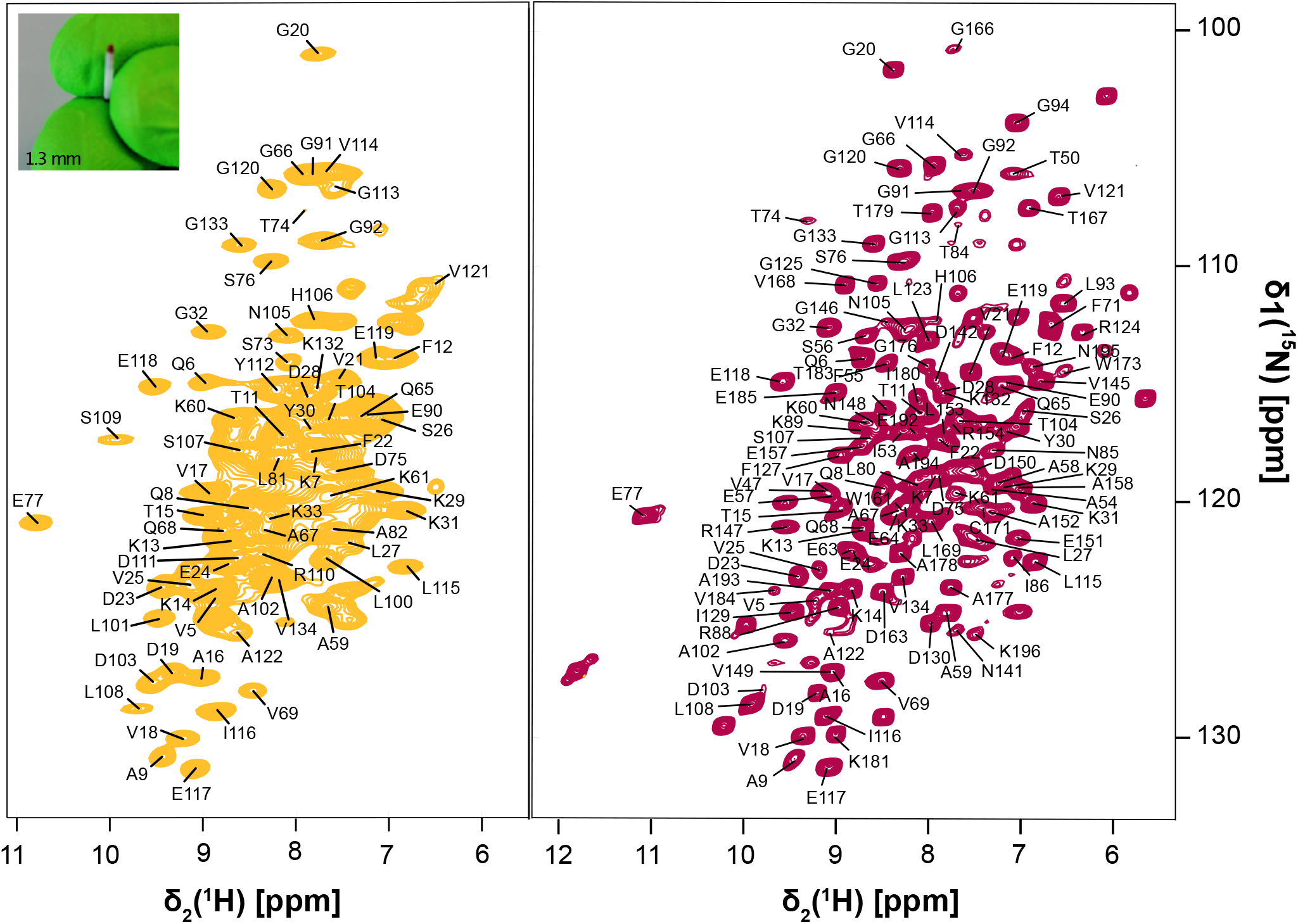
Magic-angle spinning ^1^H-^15^N correlation spectra (based on CP transfer) of Tsa1, with sequence-specific resonance assignments. Left: Tsa1^WT^ in native buffer (without any reducing agent). Right: Tsa1^WT^ in reduced state by solid state NMR.

**Fig. S4.**
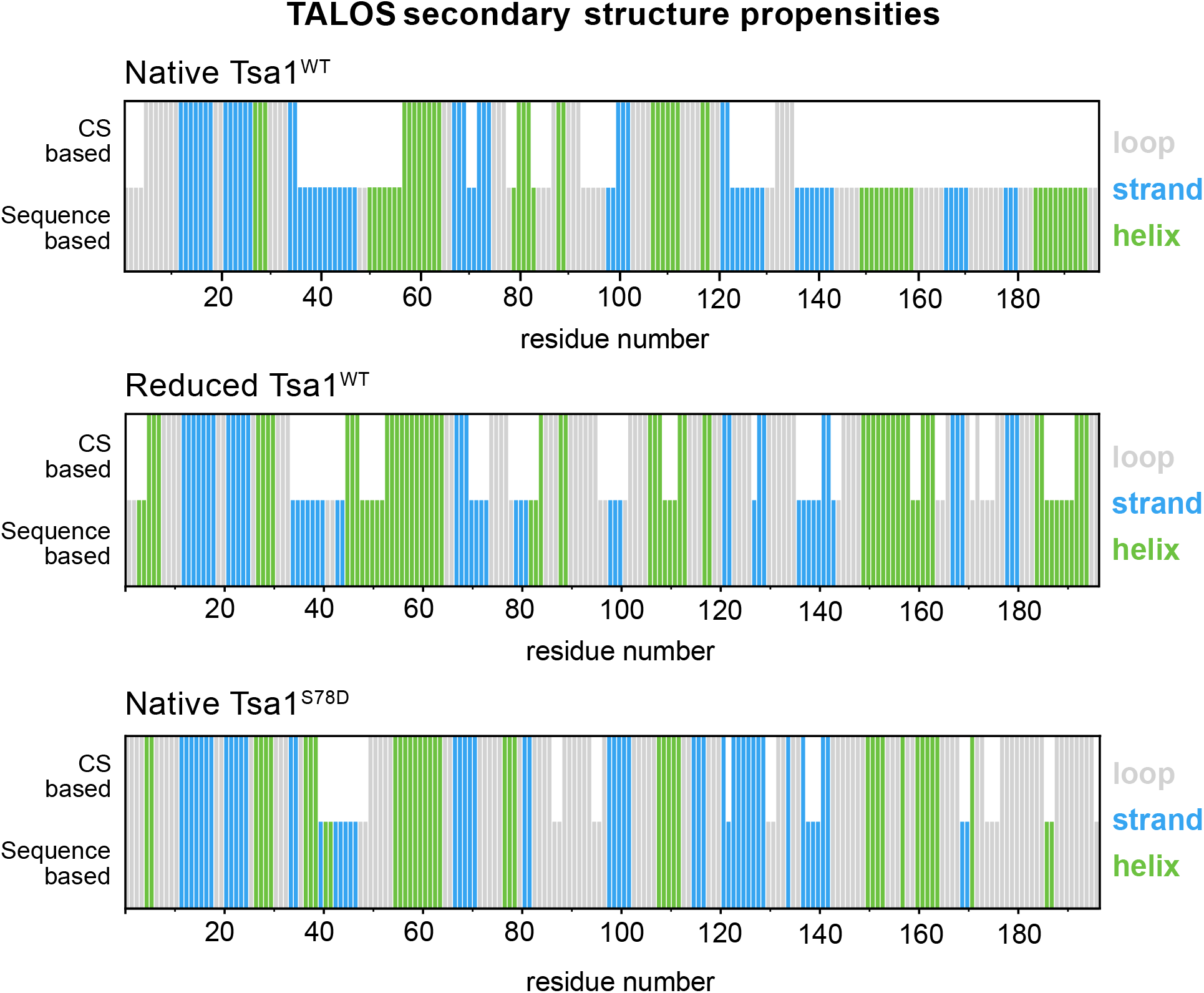
Secondary structures determined from chemical-shift assignments in decameric Tsa1 in the oxidised state (upper; corresponding to the orange spectrum in Fig. 1A; MAS NMR), decameric Tsa1 in the presence of reducing agent (middle; dark red spectrum of Fig. 1A; MAS NMR) and the dimeric mutant Tsa1^S78D^ in solution state (lower). TALOS reports secondary-structure propensities based on chemical shift of amide-^1^H, ^15^N and ^13^Cα, ^13^Cβ and ^13^CO; in the absence of chemical-shift assignments it predicts the secondary structure based on sequence (indicated by shorter bars).

**Fig. S5.**
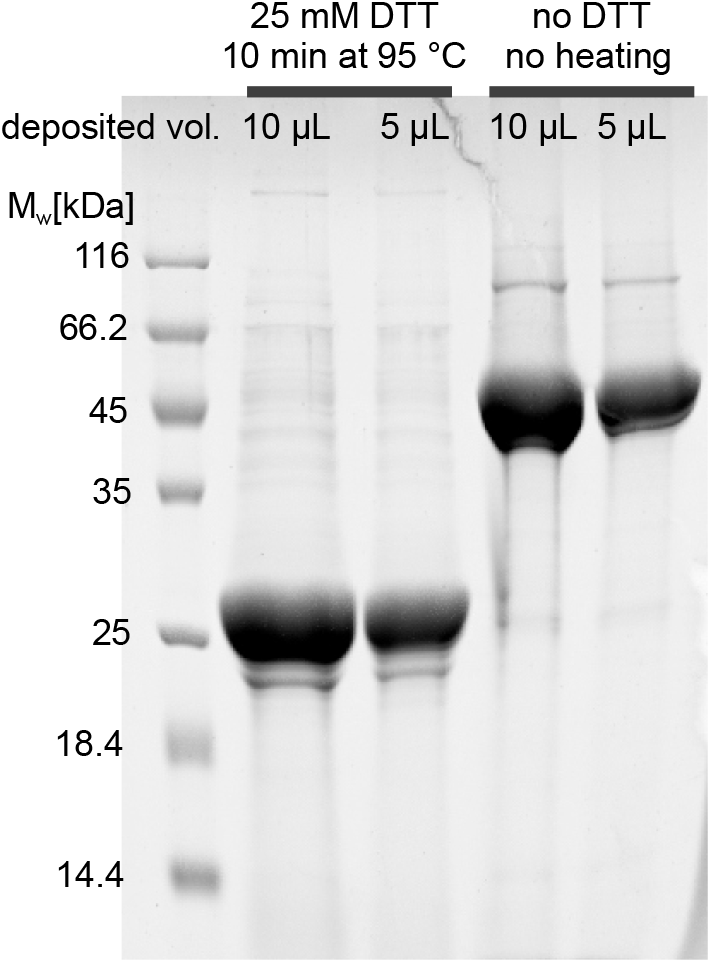
Tsa1, as obtained after purification from *E. coli* is in a disulfide-bonded state, as evidenced by SDS-PAGE analysis. Shown is the SDS-PAGE of Tsa1 that has been treated with DTT (two left lanes) or which has not been treated (right two lanes). The fact that Tsa1 migrates as a dimer without treatment and as a monomer with DTT treatment shows that the (intermolecular) disulfide bond is present in the sample obtained from purification.

**Fig. S6.**
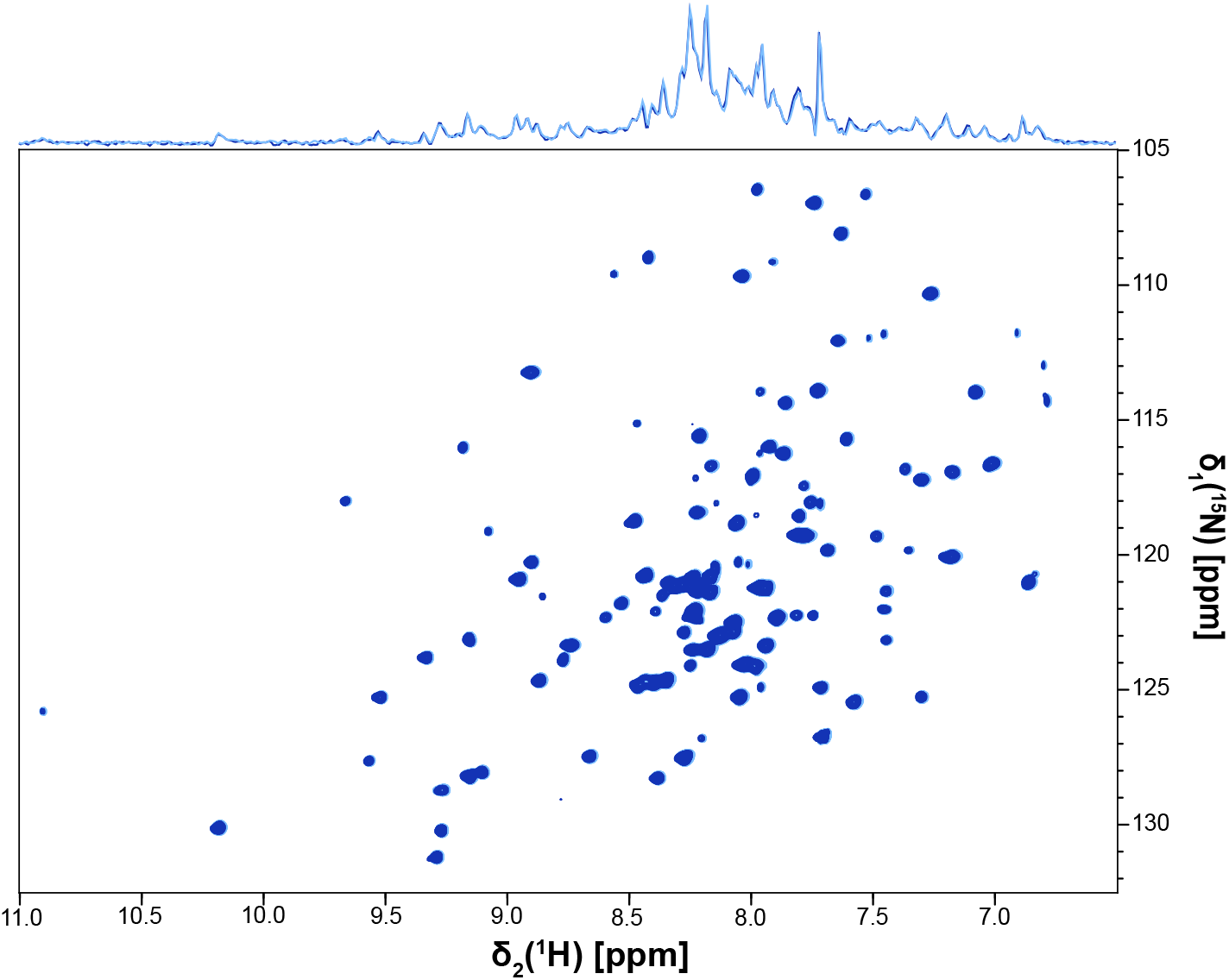
The native state obtained after expression and purification is an oxidized state. Shown are the 2D ^1^H-^15^N spectra of Tsa1^S78D^ after purification (light blue), and after incubation with 1 mM of H_2_O_2_(dark blue), as well as the 1D trace. The protein has a concentration of 500 uM in 50 mM KPi, 50 mM KCl buffer pH 7.4.

**Fig. S7.**
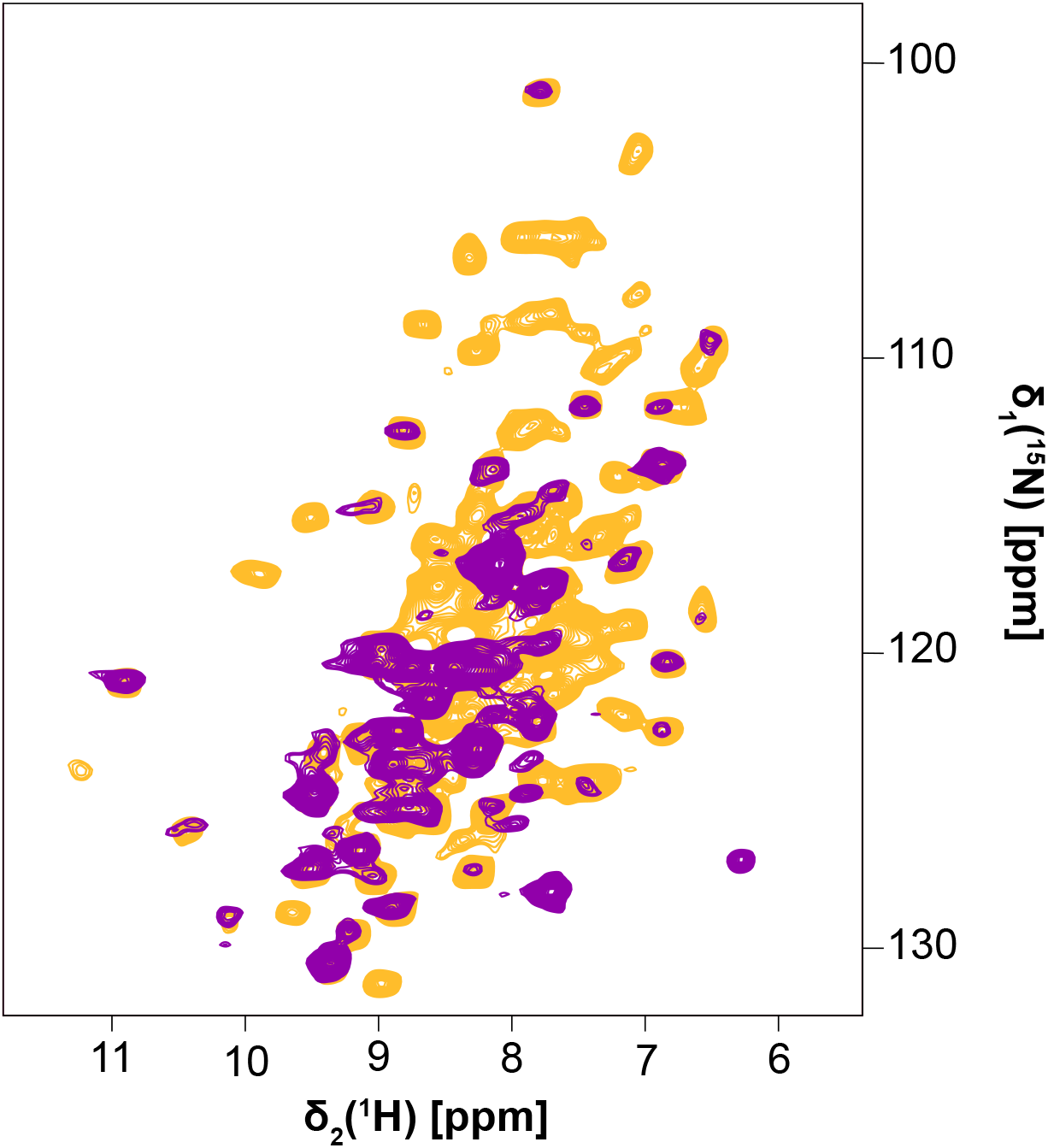
Overlay of 2D ^1^H-^15^N spectra of sedimented [U-^2^H,^13^C,^15^N] Tsa1^WT^ in 50 mM potassium phosphate, 50 mM KCl, pH 7.4, recorded with a 50 kHz MAS frequency using a CP transfer (orange) or an INEPT transfer (purple). If large-scale ps-ns motions were present, transfer with the CP scheme would be strongly reduced while INEPT transfer would be expected to be efficient; hence, peaks missing in the CP experiments may appear in the INEPT experiment. It is evident that the large number of peaks missing in the CP-based spectrum does not appear in the INEPT spectrum, suggesting that the peaks missing in the spectra of Tsa1^ox^ are missing because of μs dynamics, which broadens resonances beyond detection. This is fully in agreement with the quantitative measurements of dynamics, in particular the NERRD data. Note that there are two peaks in the INEPT spectrum which are not found in the CP spectrum, which likely are very flexible sites. We do not have assignments for these two peaks.

**Fig. S8.**
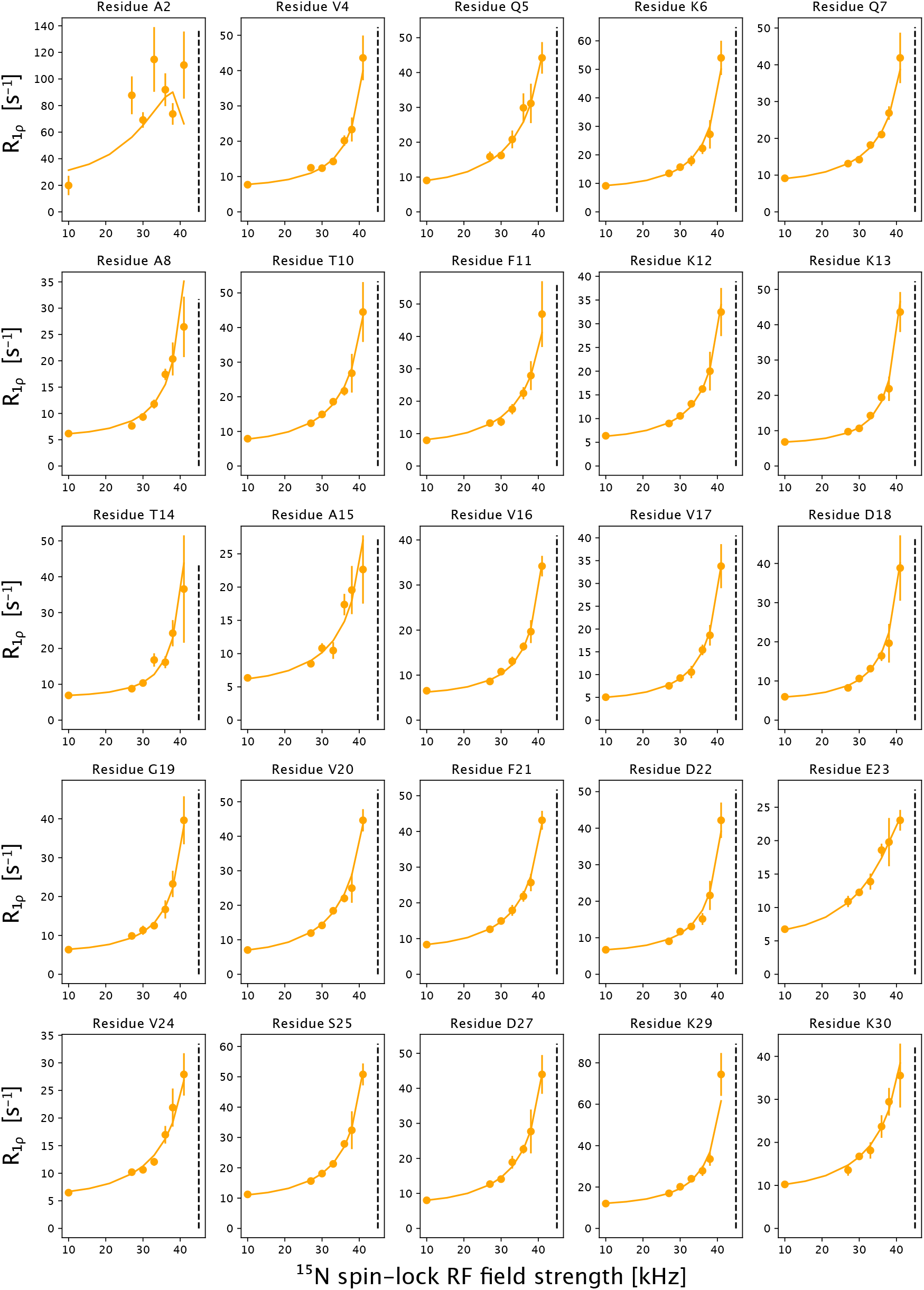
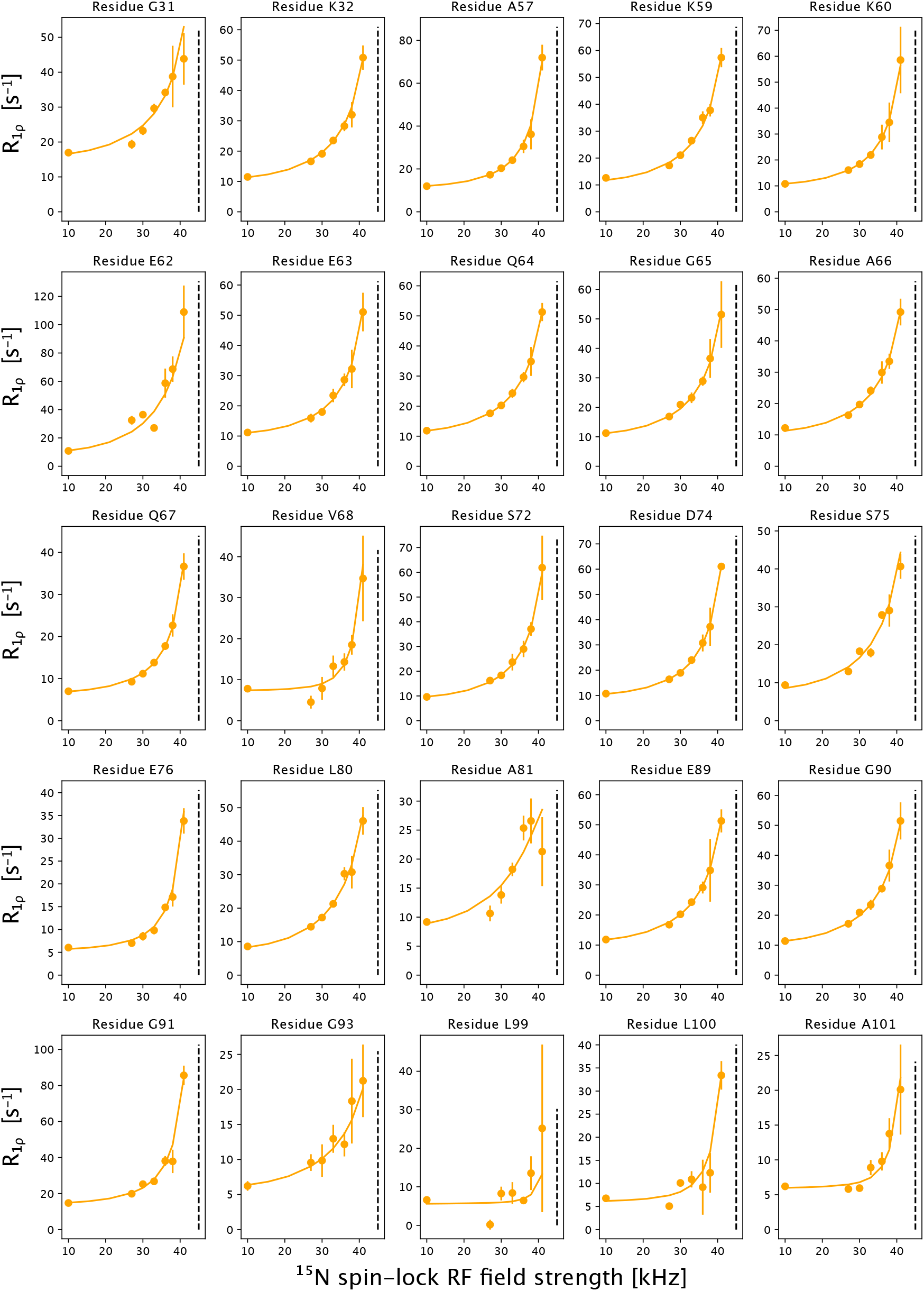
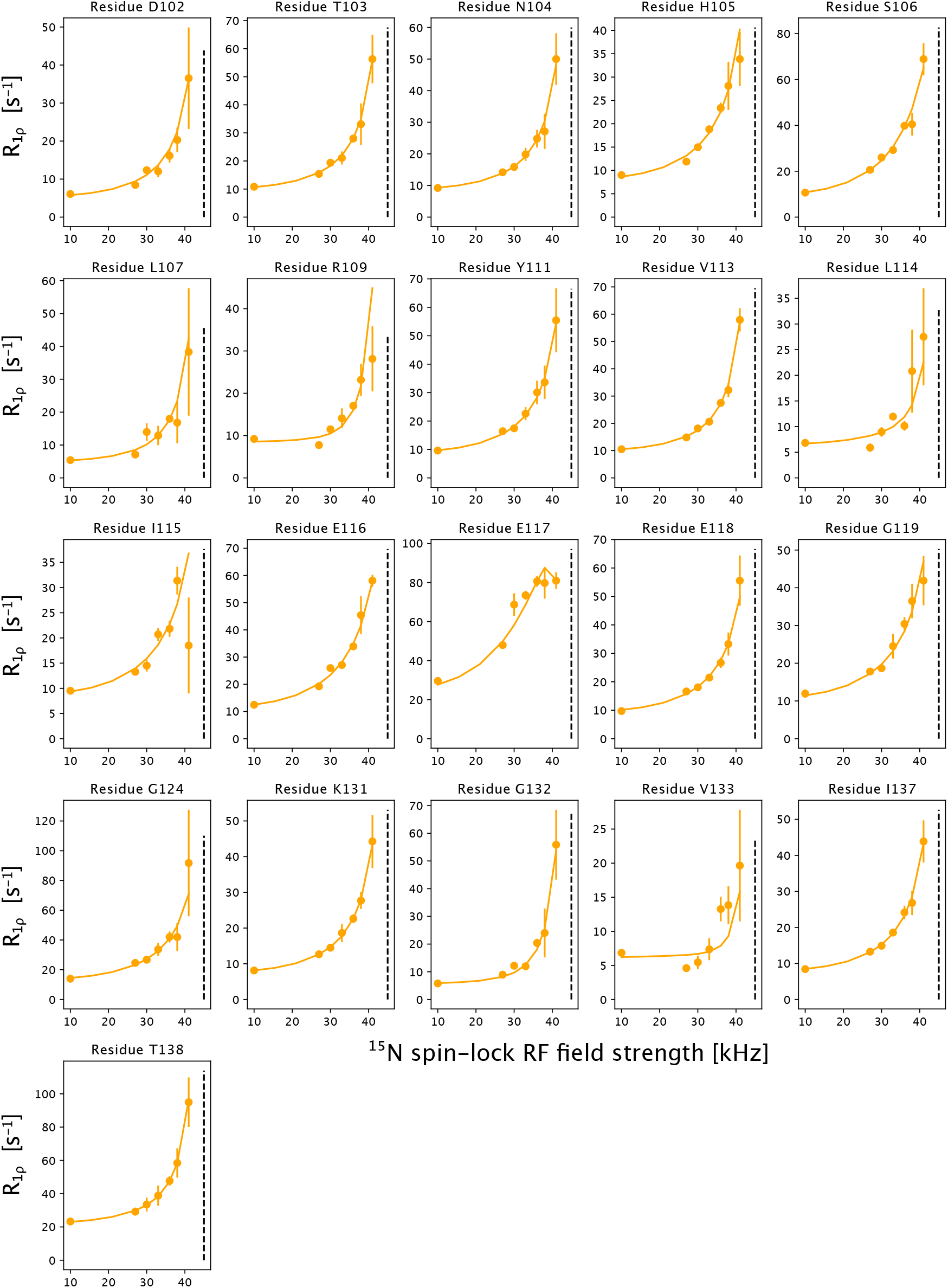
^15^N R_1ρ_ NEar-Rotary-resonance Relaxation Dispersion data of oxidised Tsa1 in the solid state. The solid line indicates the back-calculated R_1ρ_ rate constants using the DETECTORS fit with four components (see Fig. S9). The dashed vertical line indicates the n=1 rotary-resonance condition, i.e., where the RF field strength equals the MAS frequency.

**Fig. S9.**
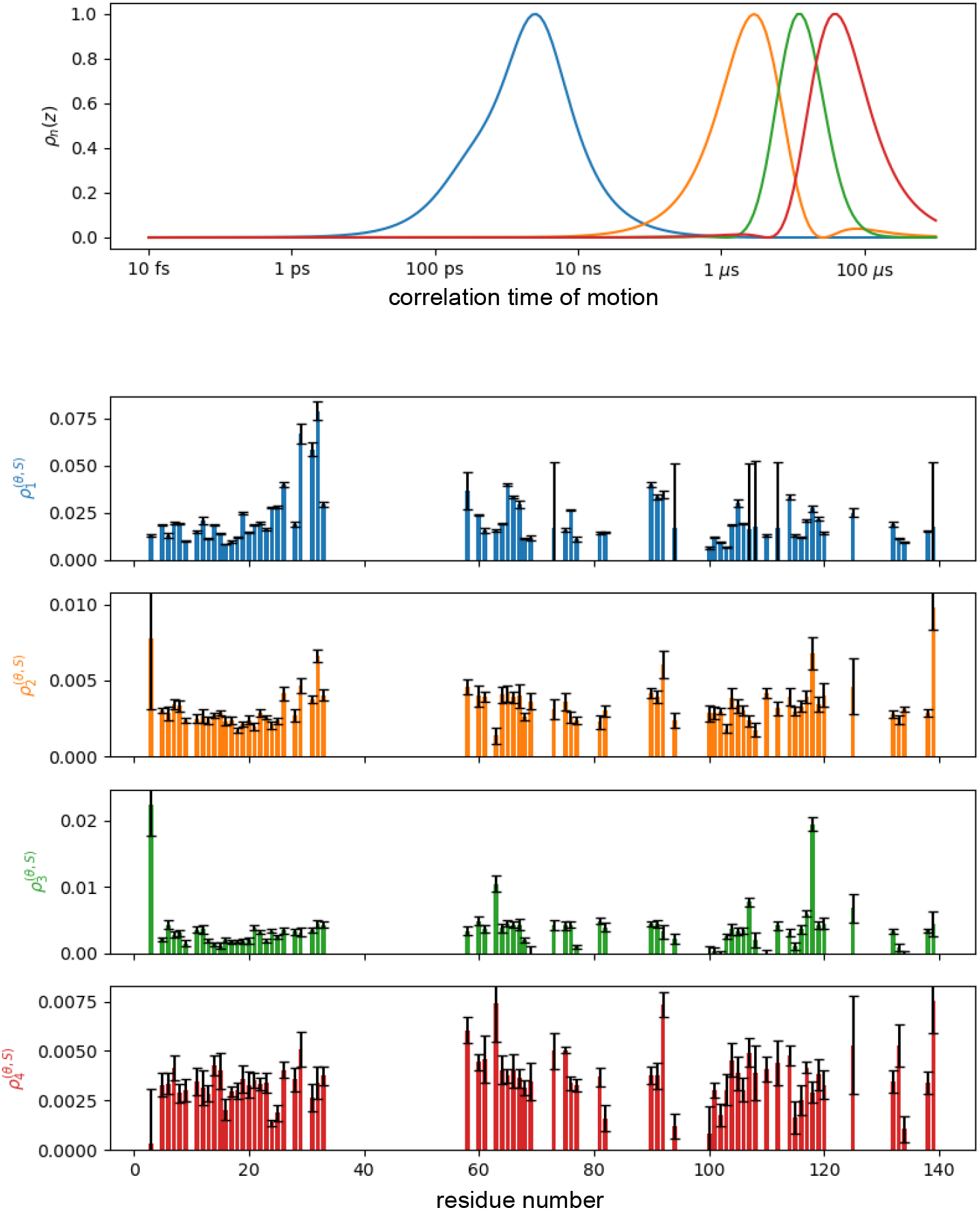
Distribution of dynamics in Tsa1^ox^ from MAS NMR relaxation data, as obtained from the DETECTORS analysis (42) (github.com/alsinmr/pyDR). All R_1ρ_ data (see Fig. S8 and the R_1_ data were used for this fit.

**Fig. S10.**
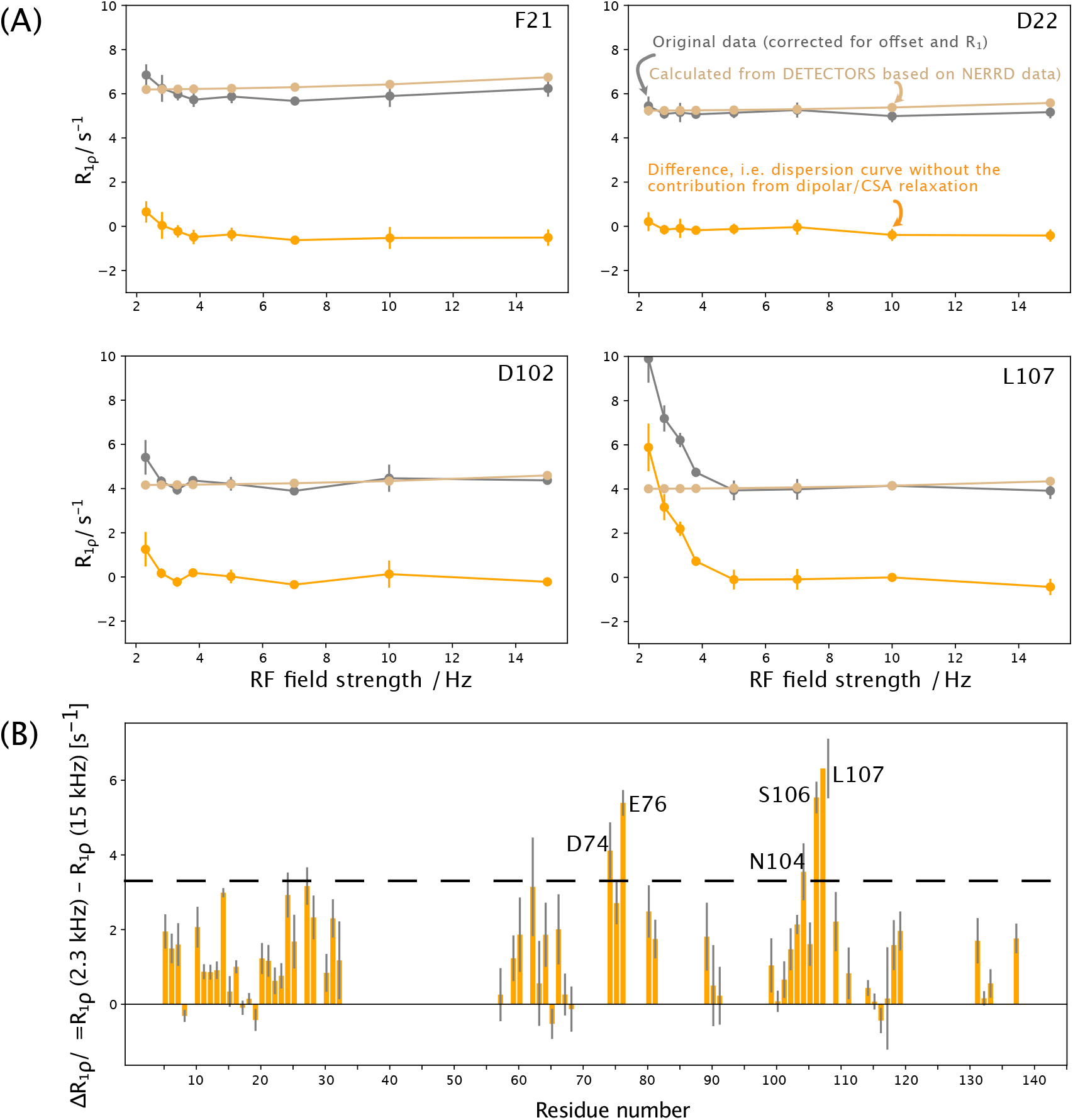
Analysis strategy and data selection of Bloch-McConnell R_1ρ_ relaxation dispersion data. (A) Outline of the procedure by which the dipolar and CSA relaxation has been accounted for in the Bloch-McConnell relaxation-dispersion profiles. The dipolar and CSA relaxation (the “tail” of the NERRD profile) contributes still to the low-RF field R_1ρ_ rate constants, which makes analysis of the chemical-shift induced relaxation dispersion complicated, and not possible with the Bloch-McConnell formalism. To account for the dipolar and CSA relaxation, we used the DETECTORS analysis with four components (see solid lines in Fig. S9) and used these fitted parameters to calculate the R_1ρ_ rate constant under the conditions at which the Bloch-McConnell RD data have been measured (55 kHz MAS, RF fields from 2.3 to 15 kHz). We subtracted these back-calculated R_1ρ_ rate constants from the experimentally measured ones. Four examples are shown. The grey data points are the original data; the brown data points are the back-calculated ones and the resulting difference is shown in orange here and also in Fig. S11. The corrected values are close to zero, showing that the DETECTORS analysis, based on R_1ρ_ rate constants measured at 45 kHz MAS and RF fields from 10 to 41 kHz, is in good agreement with these data, which had not been used in the DETECTORS analysis. Because the program used for fitting the BMRD data (relax) cannot handle negative values, we have added a uniform positive offset to these orange data, before fitting the BMRD data. (B) Difference of R_1ρ_ rate constants at the highest (15 kHz) and lowest RF field (2.3 kHz) of the data set, highlighting the residues with the most significant Bloch-McConnell relaxation-dispersion effect. In the BMRD fits, the five residues labelled in this plot have been used.

**Fig. S11.**
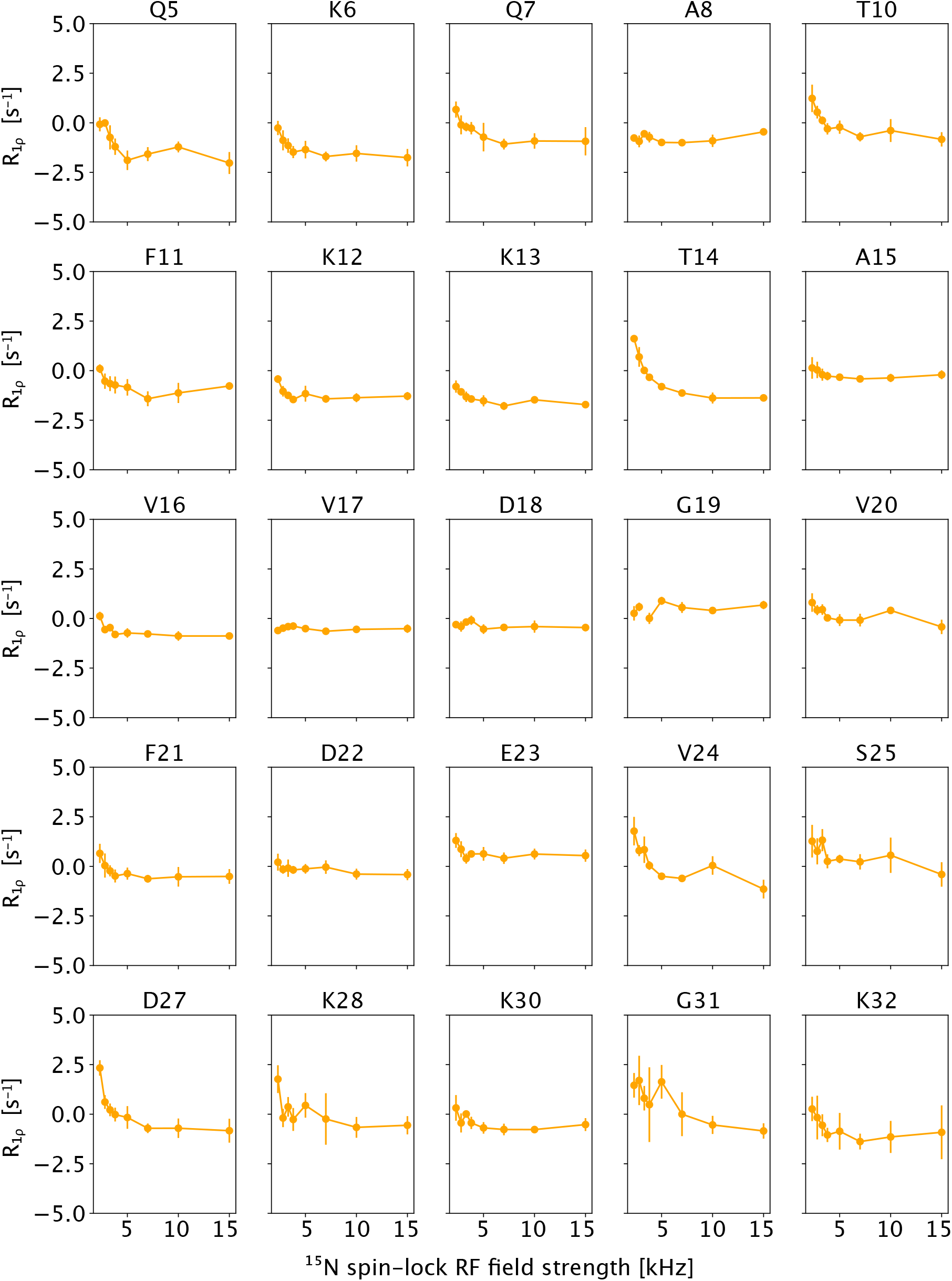
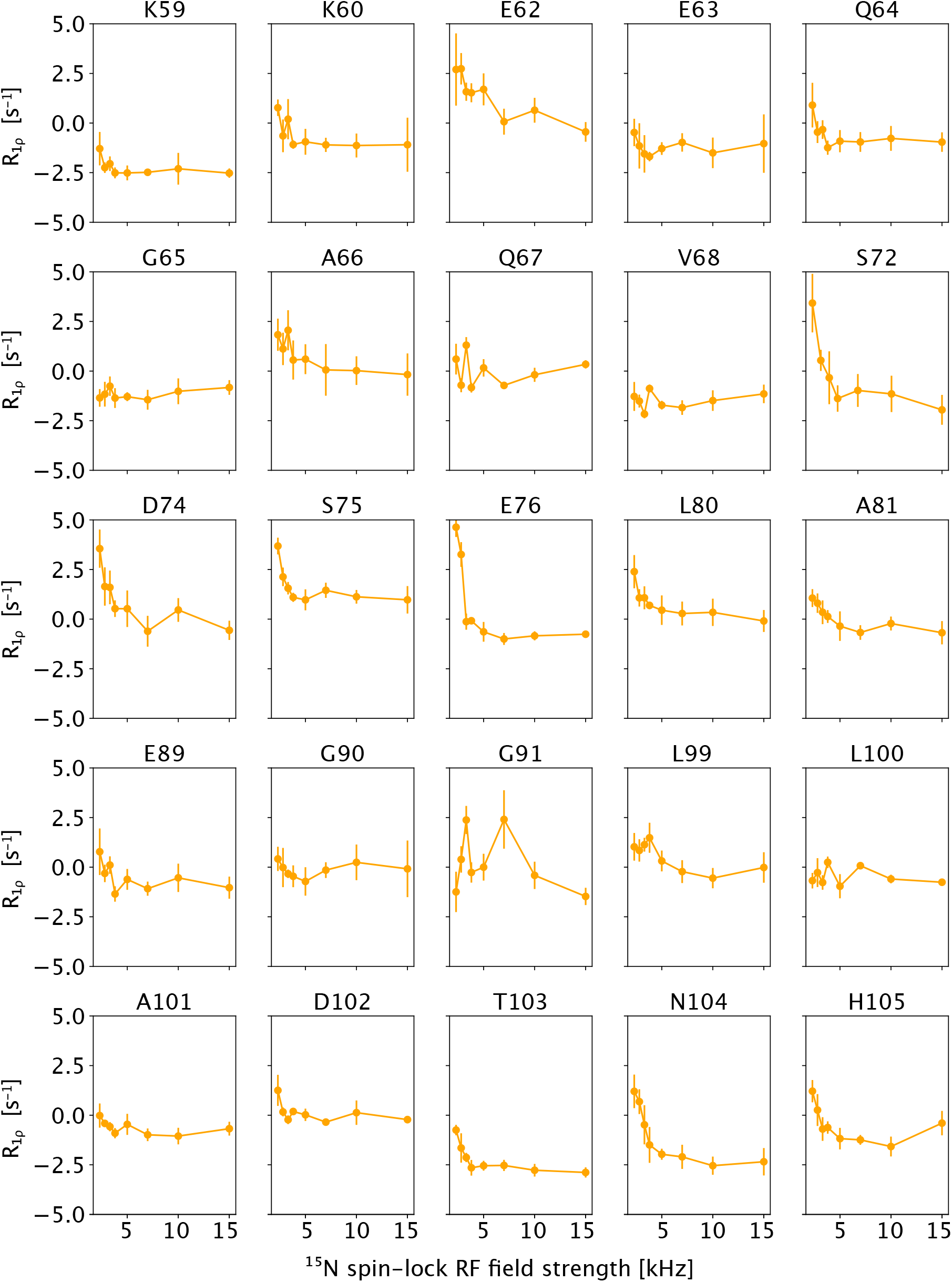
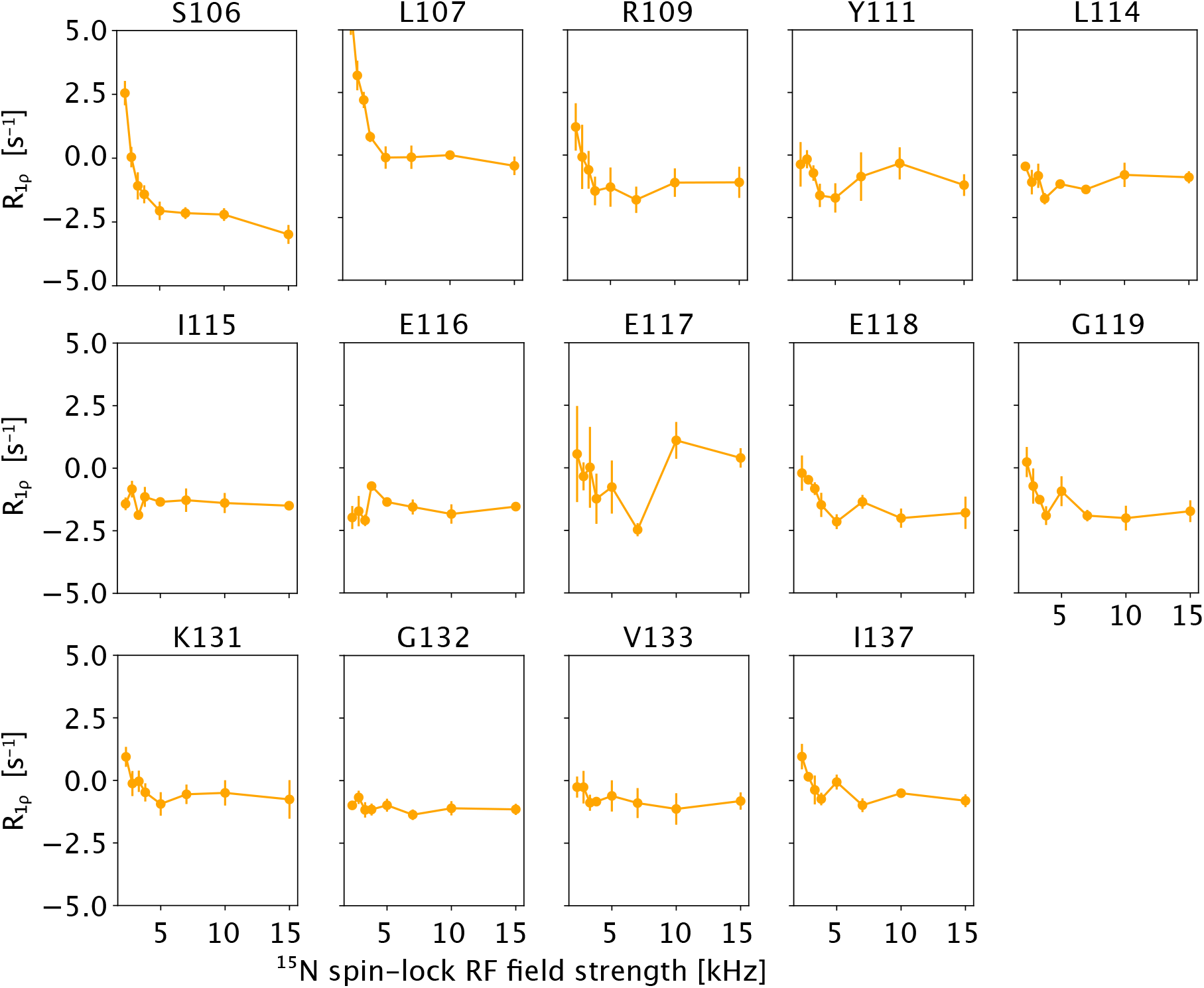
Bloch-McConnell ^15^N R_1ρ_ relaxation dispersion curves of Tsa1^ox^, corrected for resonance offset and R_1_; additionally the back-calculated rate constants based on the 4-detectors DETECTORS analysis have been subtracted from the data. See Fig. S10A for examples of this procedure.

**Fig. S12.**
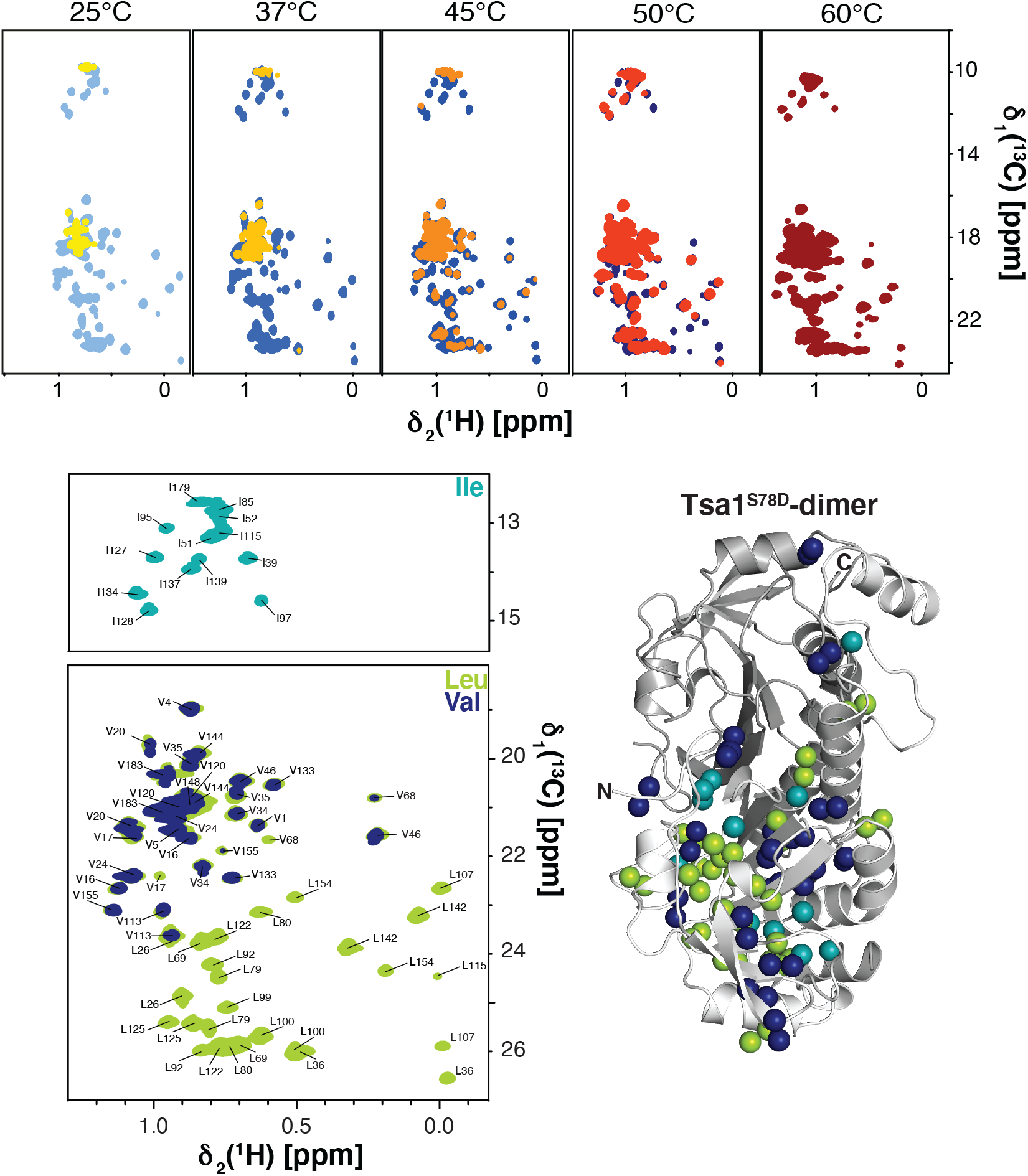
NMR of methyl labeled Tsa1. (Top) Temperature titration of [U-^2^H,^15^N, Ile-δ1-^13^CH_3_, Leu/Val-^13^CH_3_] labeled Tsa1^WT^ (yellow to red) and [U-^2^H,^15^N, Ile-δ1-^13^CH_3_, Leu/ Val-^13^CH_3_] labeled Tsa1^S78D^ (light blue to dark blue) by solution state NMR, ranging from 25°C to 60°C. Data show that Tsa1^S78D^ is not impacted by temperature change. Therefore its structure is conserved. In contrast, for Tsa1^WT^, the spectrum only shows signals of few residues at 25° C. These peaks tend to correspond to the most flexible parts of the protein, such as I179, I85 or V183, for which the order parameters are low (see Fig. 4); the large-amplitude motion of these sites increases their coherence life times and it is, thus, expected that they are visible even at low temperature. When increasing the temperature one retrieves the same NMR spectrum as the one obtained for the dimeric mutant. This is understandable as high temperature promotes the dissociation to dimers (see SEC-MALS data in Fig. S1). (Bottom) 2D ^1^H-^13^C spectrum of [U-^2^H,^15^N, Ile-δ1-^13^CH_3_, Leu/ Val-^13^CH_3_] Tsa1^S78D^, with manual assignment achieved using ^13^C-methyl-SOFAST-NOESY experiments with mixing times of 50 ms and 600 ms as well as a 3D HMBC-HMQC experiment. Spectra from two different samples are shown in the lower panel, from either labeling Val and Leu simultaneously (green) or from labeling only Val (blue). The right panel shows the location of Ile, Leu and Val methyl groups on a monomeric subunit of the Tsa1 structure.

**Fig. S13.**
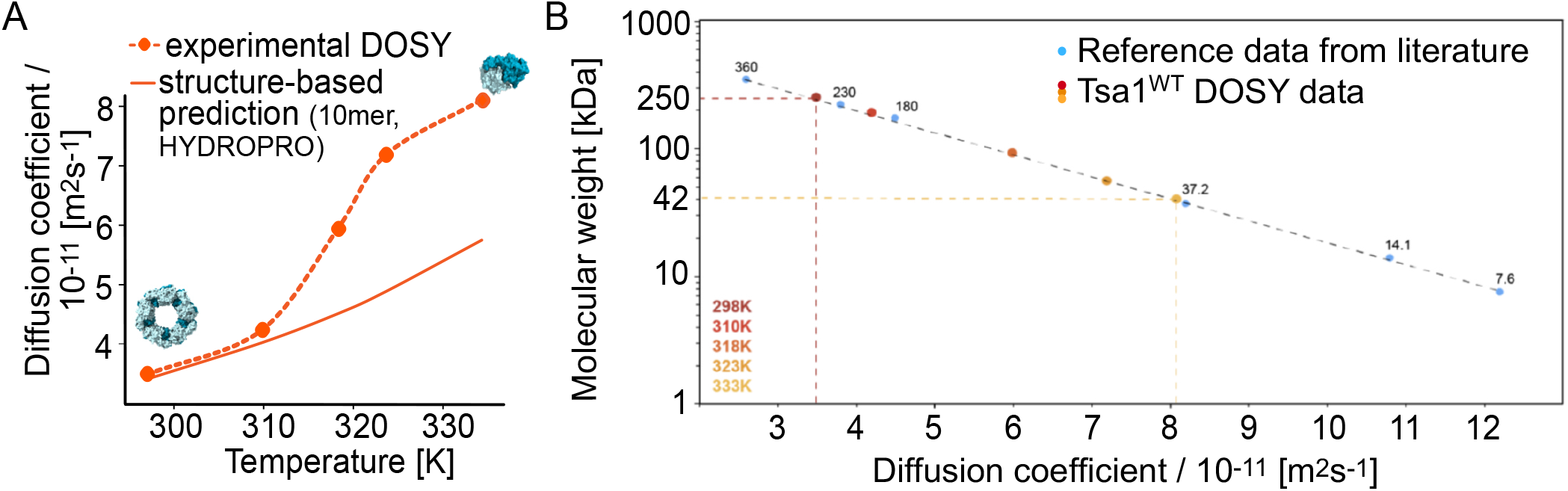
Diffusion-ordered spectroscopy (DOSY) data measured by methyl-directed experiments of wild-type Tsa1. Left: fitted diffusion coefficients as a function of temperature. The solid orange line is the predicted diffusion coefficient using the program HYDROPRO (76). Right: estimated molecular weights. To construct the calibration curve on the right, the following published translational diffusion coefficients, measured at 25°C, were used: immunoglobin-binding domain of streptococcal protein G, GB1 (MW = 6.2 kDa) (75), lysozyme (MW = 14.1 kDa) and interleukin-10 (MW = 37.2 kDa) (74), bacterial HsIV (MW = 230 kDa), one-half proteasome from *Thermoplasma acidophilum*, α7α7 (MW = 360 kDa), and the α7 single ring variant of the proteasome (MW = 180 kDa) (73).

**Fig. S14.**
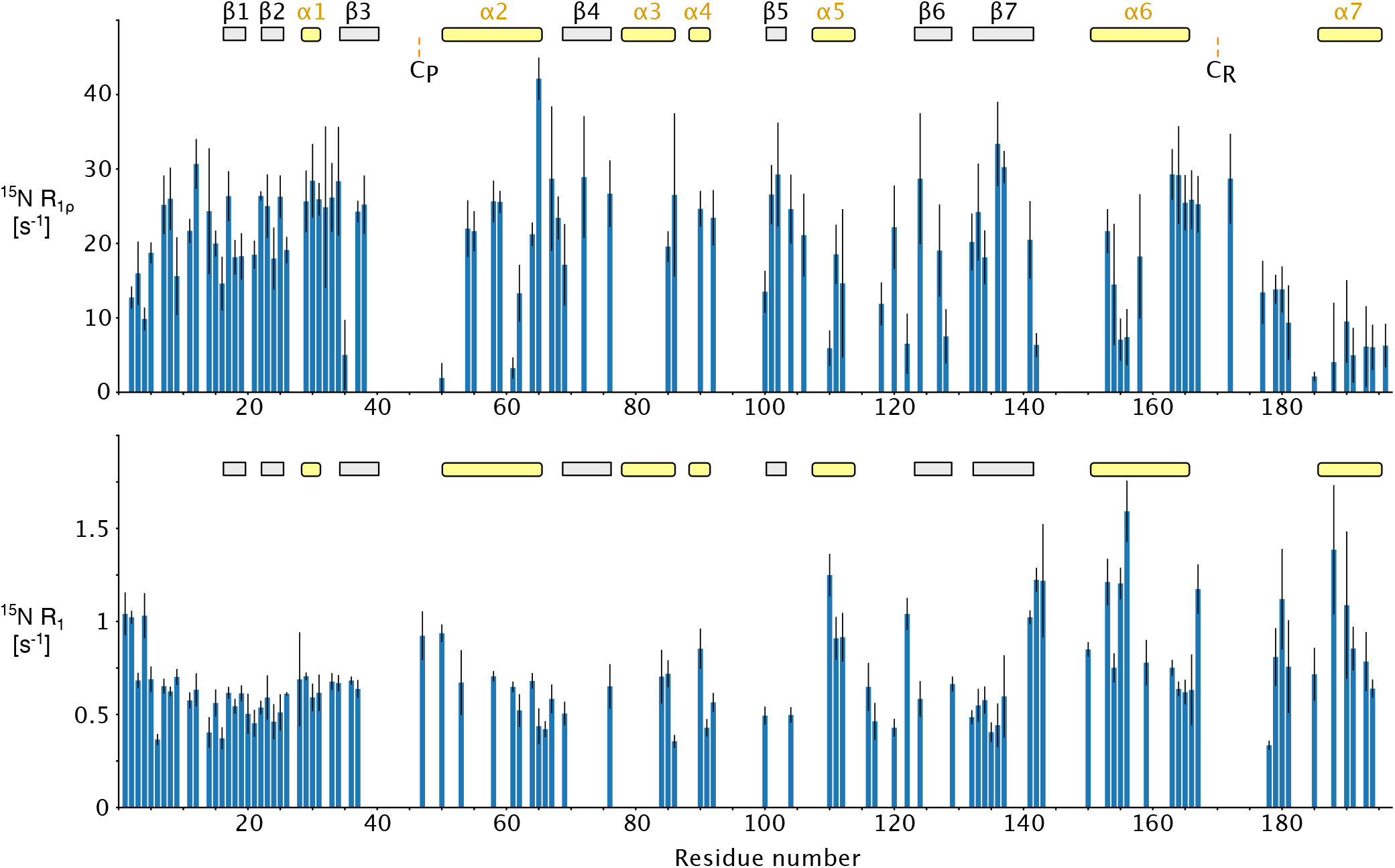
^15^N spin relaxation data of the dimeric Tsa1^S78D^ mutant at 37 °C. The indicated secondary structure elements above the plots are obtained from the decameric state (PDB 3SBC). It is evident that the helix α7 has higher flexibility than the other secondary structure elements. Together with the TALOS secondary-structure data (Fig. 1), this observations shows that in the dimer the C-terminal helix is disordered.

**Fig. S15.**
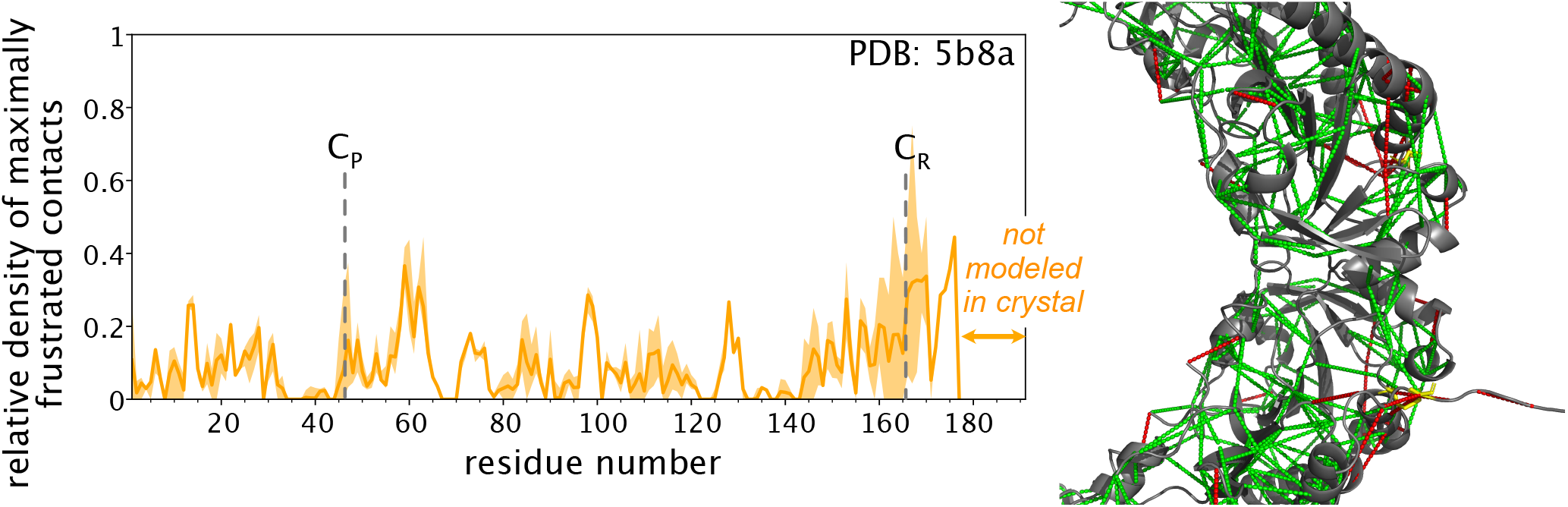
Structural frustration data of the oxidised PRDX from *E. coli*, PDB entry 5B8A.

**Fig. S16.**
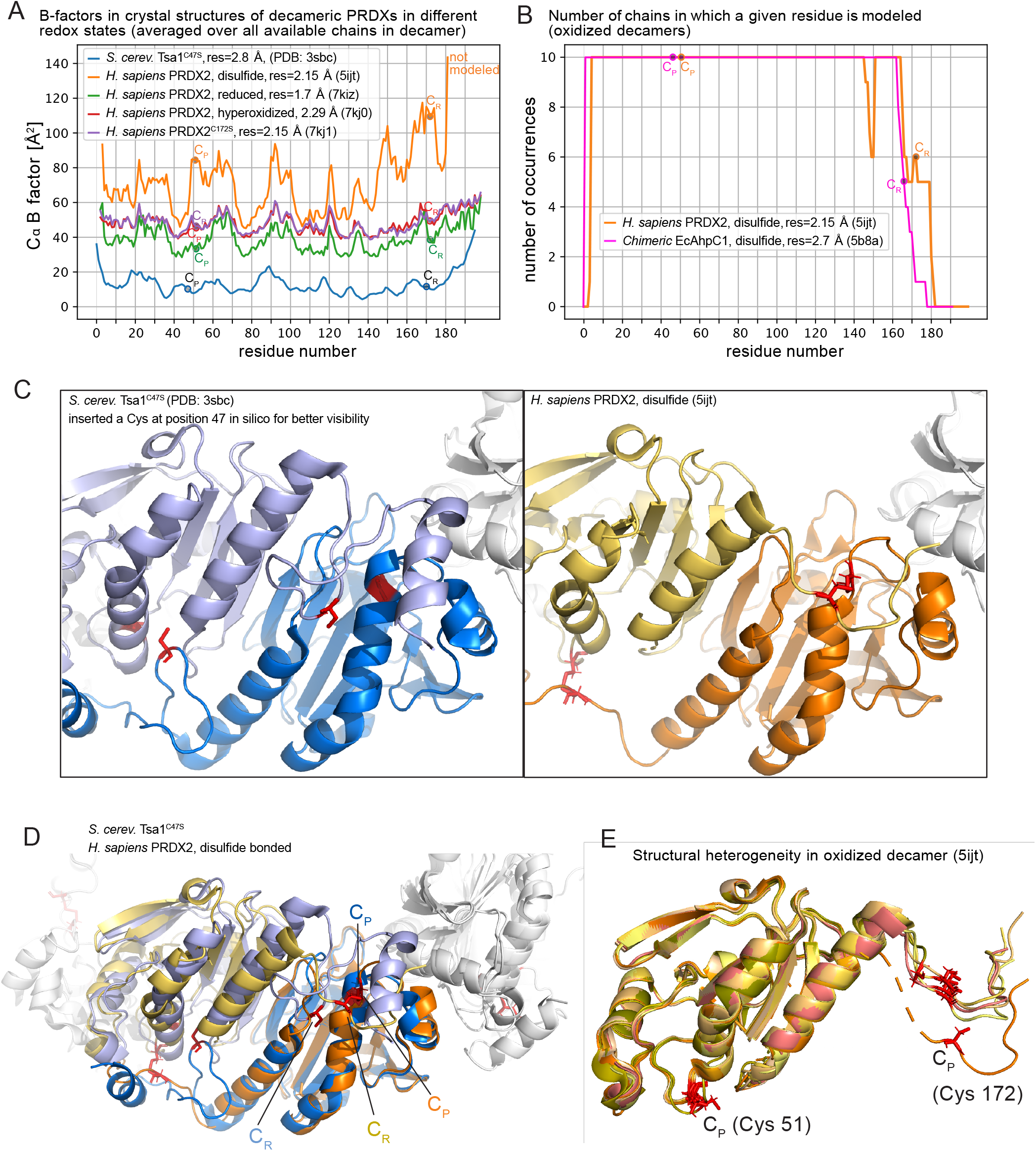
Insights into PRDXs structure from crystallography. A) B-factors of decameric PRDXs Cαin different oxidation states. B) Occurences of modelized residues over the sequence of the oxidized decamer of hPRDX2, showing that the C-terminal part of the protein cannot be properly modeled because of it intrinsic dynamics or lack of structure. C) Comparison of the α2 helix structure between reduced *Sc* Tsa1 (3SBC) with insertion of the wild type Cys47 for better visibility, and oxidized *Hs* PRDX2 (5IJT) in its S-S state. D) Overlay of figures C for better understanding. E) Structural ensembles of oxidized decamer of *Hs* PRDX2 (5IJT) showing structural heterogeneity of the C-terminal tail.

**Fig. S17.**
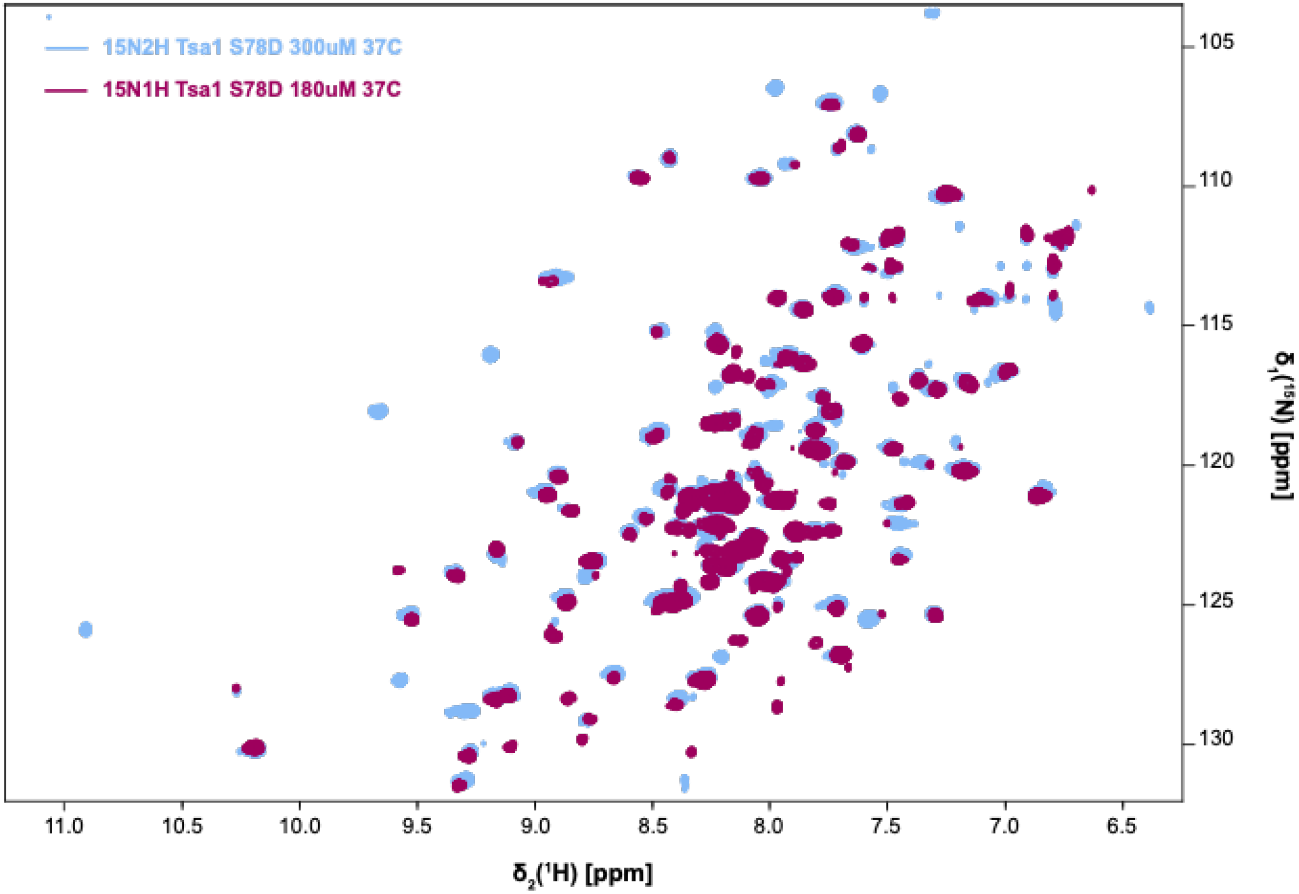
Comparison of [^15^N-^1^H] TROSY HSQC solution-NMR spectra of a deuterated sample, produced in D_2_O-based growth medium ([U-^2^H,^15^N] Tsa1^S78D^, light blue), and a protonated sample ([U-^15^N] Tsa1^S78D^, red).

